# Biomass-derived Lignin Nanoparticles for the Sustained Delivery of Vascular Endothelial Growth Factor-C

**DOI:** 10.1101/2025.04.23.649697

**Authors:** Ahmad Mavali Zadeh, Emanuela Gatto, Raffaella Lettieri, Honey Bokharaie, Alice Caravella, Cadia D’Ottavi, Elisabetta di Bartolomeo, Fabio Domenici, Setareh Sima, Alexandra Correia, Khushbu Rauniyar, Anna Klose, Zahra Gounani, Timo Laaksonen, Jaana Künnapuu, Michael Jeltsch

## Abstract

Vascular Endothelial Growth Factor C (VEGFC) is a promising biological drug, as preclinical studies have shown its potential in treating a variety of conditions, including myocardial infarction and neurodegenerative diseases. For lymphedema, a disease that currently can only be treated symptomatically, adenoviral VEGFC gene therapy has been evaluated up to phase II studies. However, the AdVEGFC is rapidly inactivated by the immune system, and alternative delivery methods might yield better results. Thus, we wanted to investigate the synthesis, characterization, and stability of lignin nanoparticles (LNPs) as carriers for VEGFC.

As biomass-derived lignin nanoparticles provide a sustainable, cost-effective, and tunable platform for drug delivery, with the potential to enhance drug stability and release, lignin was extracted from wood biomass derived from grape shoots using the organosolv method and subsequently synthesized into nanoparticles. The resulting lignin nanoparticles (LNPs), with an average size of 142±62 nm and a zeta potential of -40±8 mV, were characterized through comprehensive techniques, including spectroscopy and microscopy, to gain insights into their structural and morphological properties.

Furthermore, the loading and release efficiency of VEGFC onto LNPs were evaluated, demonstrating effective loading and controlled release. Stability tests in plasma and cell proliferation/viability assessments using the MTT (3-[4,5-dimethylthiazol-2-yl]-2,5 diphenyl tetrazolium bromide) assay were conducted to assess biocompatibility and therapeutic potential. Our data shows that VEGFC is a relatively stable protein and that the primary advantage of nanoparticle-based delivery would be to delay release as opposed to protect VEGFC from degradation/inactivation.

**Graphical Abstract:** Flow chart of the generation and analysis of biomass-derived VEGFC-loaded lignin nanoparticles.

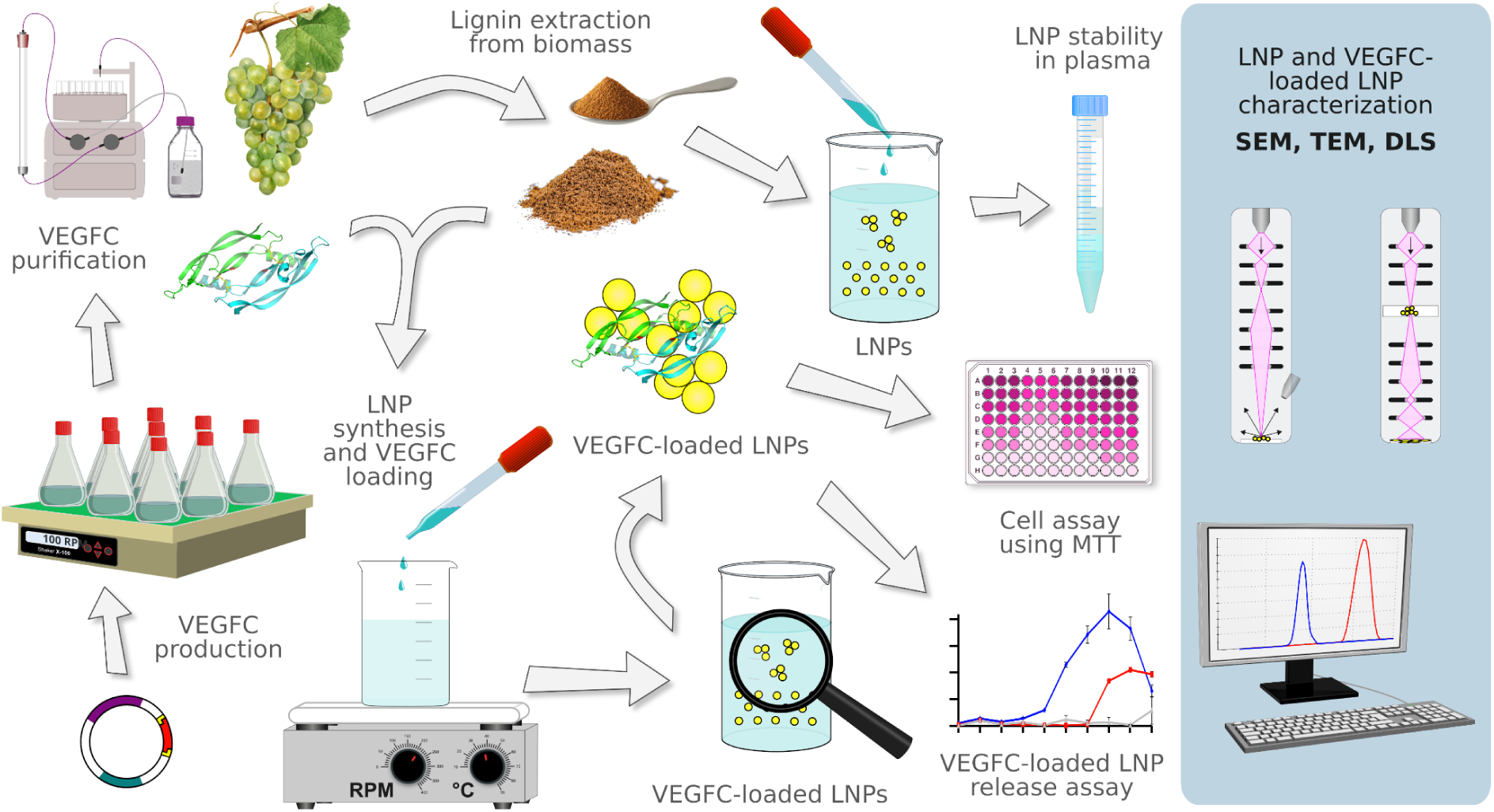

## 1. Introduction

### Lignocellulosic Biomass and Lignin Structure

Lignocellulosic biomass, a readily available resource here on Earth, carries enormous potential that exceeds its current utilization.^1^ Lignin, an important component of lignocellulosic biomass, is a versatile biopolymer found abundantly in plant cell walls. It is the second most abundant polymer on Earth.^2^ Lignin has emerged as a remarkable material with a wide range of applications across various fields.^3^ However, lignin is a structurally complex aromatic polymer highly resistant to chemical and biological breakdown, and it remains challenging to utilize lignin fully.^4^

Each type of lignocellulosic biomass comprises varying proportions of lignin, hemicellulose, and cellulose, which influence its properties.^5^ Lignin was extracted from wood biomass derived from grape shoots, an abundant agricultural byproduct of vineyard pruning. Utilizing grape shoots as a lignin source promotes sustainability by adding value to waste materials while reducing environmental impact. According to Jimenez et al. Grape shoot contains approximately 20% lignin.^6^ Lignin is characterized by its irregular and random cross-linking. It consists of three primary phenylpropane monomers: *p-*coumaryl alcohol (H unit), coniferyl alcohol (G unit), and sinapyl alcohol (S unit). These units differ mainly in the number of methoxy groups attached to their aromatic rings.

The heterogeneity of lignin arises from the interaction between its H, G, and S units, which creates various functional groups and linkages.^7^ According to the literature, seven predominant linkages are β–O–4–aryl ether (β–O–4), 4–O–5–diaryl ether (4–O–5), β–5–phenylcoumaran (β–5),5–5–biphenyl (5–5), β–1–(1,2–diarylpropane) (β–1) and β–β (resinol) in the lignin molecules.^8,9^ The aryl ether β-O-4 bond is the most prevalent, comprising about 50% of the linkages.^10^ Consequently, the composition and characteristics of lignocellulosic biomass are crucial factors in determining the suitable pretreatment method. The primary objective of pretreatment is to disassemble ester and ether bonds within lignocellulosic biomass, allowing for the isolation of its pure components.^11^ Various pretreatment methods have been extensively researched and documented in numerous review articles over the years.^11,13^ Among these, organosolv pretreatment is particularly notable for its ability to separate each biomass component effectively. This method yields relatively pure lignin, which can be sold as a by-product or transformed into higher-value products within a biorefinery framework.^15^

### Lignin Nanoparticles (LNPs)

Lignin-derived nanoparticles (LNPs) show great promise in various biomedical applications due to their unique properties and low cytotoxicity. Dai et al. (2017) developed LNPs assembled with resveratrol (RSV) and Fe_3_O_4_ magnetic nanoparticles, which demonstrated good anticancer performance, controlled RSV release, and stability, without causing harmful effects on cells or mice.^16^ Similarly, Figueiredo et al. (2017) created both pure and iron-infused LNPs, finding them effective in drug delivery against cancer cell proliferation with low cytotoxicity and hemolytic rates (10–12%) after 12 hours of incubation.^17^

Pourmoazzen et al. (2020) formulated cholesterol-modified LNPs for pH-responsive drug delivery, achieving a 55% release of folic acid at pH 8.0.^18^ Zikeli et al. (2020) prepared LNPs with entrapped essential oils for biocide delivery, showing a slower release compared to non-entrapped oils, which is beneficial for wood preservation.^19^ Tortora et al. (2014) developed oil-filled lignin microcapsules for storing and delivering hydrophobic molecules. They successfully incorporated and released Coumarin-6 in Chinese hamster ovary cells without toxicity.^20^ The safety and efficacy of LNPs in various applications, such as drug delivery, biocide delivery, and gene therapy, have been demonstrated across multiple studies, indicating their potential for future biomedical applications.^21,22^

### Vascular Endothelial Growth Factor-C (VEGFC)

VEGFC is one of the two essential members of the VEGF family. VEGFs are the primary growth factors for angiogenesis and lymphangiogenesis. Angiogenesis, the formation of new blood vessels from pre-existing ones, occurs under physiological circumstances such as wound healing, organ regeneration, embryonic development, the formation of collateral blood vessels in response to ischemia, and female reproductive processes like ovulation and placenta formation. However, VEGFs are also responsible for the pathologic angiogenesis in many diseases, including cancer, rheumatoid arthritis, diabetic retinopathy, and psoriasis.^23^

In pancreatic endocrine tumors, as well as in MCF-7 breast cancer and β-cell tumor models, the upregulation of VEGFC has been linked to progression and metastasis, demonstrating its role in tumor biology and potential as a therapeutic target.^24,25,26,27^ VEGFC might also explain the prognostic value of prostate-specific antigen (PSA) in prostate cancer, as PSA was shown to convert pro-VEGF-C into active VEGFC,^28^ and VEGFC neutralization with sozinibercept is being tested at the moment in two phase III clinical trials (<u>NCT04757610</u>, <u>NCT04757636</u>). Moreover, pro-angiogenic therapy with VEGF or VEGFC presents a therapeutic option for neuronal and cardiovascular diseases, extending their role beyond an anti-angiogenic drug target in cancer and ocular neovascularization. Preclinical research indicates that VEGFC is crucial in neurogenesis, aids in coronary artery development, and contributes to cardiac lymphatic repair.^29,30,31^ Delivering growth factors effectively and safely is challenging primarily due to the difficulty of achieving a physiological distribution across the target tissues. Moreover, maintaining therapeutic levels poses a challenge, as frequent administration, such as daily injections, can be prohibitively expensive.

The encapsulation of VEGFC in nanoparticles offers several benefits that address some of these challenges. Loading VEGFC onto nanoparticles could significantly enhance its stability and protect it from premature degradation. It also allows for active targeted delivery to specific sites, such as tumor tissues, via nanoparticle modification, e.g., using ligands or antibodies binding to cell surface receptors that target cells express in high amounts. Thus, the local concentration of the nanoparticle payload increases, and off-target effects are reduced.^32^ E.g., decoration with hyaluronic acid could increase the drug concentration in the lymphatic system via interaction with the lymphatic-specific hyaluronic acid receptor LYVE-1,^33^ enhancing its therapeutic efficacy while reducing potential side effects associated with systemic distribution.

VEGFC has been used, for example, in the experimental treatment of breast cancer-associated lymphedema. An adenoviral vector was designed to provide VEGFC for a limited time to facilitate the integration of transplanted lymph nodes into the local lymphatic network.^34^ In a similar fashion, VEGF-C-charged nanoparticles could be fine-tuned to release the growth factor in a controlled manner, providing a sustained therapeutic effect. The combination of enhanced stability, targeted delivery, and controlled release presents a compelling case for using polymeric nanoparticles to advance the therapeutic applications of VEGFC. This strategy has the potential to overcome some of the barriers that have limited the clinical use of growth factors and to open new ways to treat diseases that involve vascular growth and repair.^35^

This study investigates the possibility of using LNPs as carriers for VEGFC. The research encompasses several objectives. Firstly, this study explores the distinctive characteristics of LNPs, using waste biomass as a sustainable source for their preparation. Secondly, the study evaluates the efficiency of loading VEGFC onto LNPs and examines their release profiles under physiological conditions. Biocompatibility is assessed by evaluating the stability of LNPs in plasma and performing cell proliferation/viability assays, as both contribute to a comprehensive biocompatibility assessment.

## 2. Experimental Section

### 2.1. Lignin Extraction

Lignin was extracted from grapevine shoot biomass collected from a southern Italy vineyard. 4 g of powdered sample, which had already been dried in an oven at 60°C for 2 hours, was mixed with a formic acid/acetic acid/water solvent (30/50/20 v/v/v). The ratio of solid to liquid was 1:25. The mixture was placed in a round-bottom flask, and the reaction conditions were set to 107°C and 3 hours.^36^ After extracting, the yield of lignin was determined according to the formula *yield (%) = (g of extracted lignin) / (g of attributed lignin content)* ×*100.*^6^

### 2.2. Characterization of Lignin

Lignin was characterized using several analytical techniques: Fourier Transform Infrared Spectroscopy (FTIR) for analyzing chemical composition and structural changes; ultraviolet-visible spectroscopy (UV-Vis spectroscopy) to investigate its electronic structure; Differential Scanning Calorimetry (DSC) to assess thermal properties, including thermal stability; Thermogravimetric Analysis (TGA) for evaluating thermal stability and decomposition behavior; and X-ray Diffraction (XRD) to examine crystalline structure and confirm the amorphous nature of the samples.

A Thermo Scientific Nicolet iS50 FT-IR Spectrometer in ATR mode equipped with OMNIC software was used to conduct the FT-IR analysis. The experimental parameters were set to a wavenumber range of 4000-600 cm⁻¹, a resolution of 4 cm⁻¹, and 32 scans per sample. For background measurements, 64 scans were recorded prior to each sample analysis. The resulting spectra were subsequently processed and normalized using GraphPad Prism 8 software, with normalization referenced to the peak at 1025 cm⁻¹.

UV-Vis spectroscopy was performed using a CARY 100 Scan UV-VIS Spectrophotometer. The lignin samples were dissolved in a 0.01 M sodium hydroxide solution to achieve a concentration of 0.1 mg/ml. The spectra were recorded over a wavelength range of 200-800 nm with a resolution of 1 nm and a scan speed of 200 nm/min. A quartz cuvette with a path length of 1 cm was used. The baseline was corrected by subtracting the solvent’s absorbance spectrum from the sample spectra.

Simultaneous Thermogravimetric Analysis and Differential Scanning Calorimetry (TG-DSC) in air were performed using a Mettler Toledo TG-DSC 1, Star System. Samples were placed in a platinum pan and heated from room temperature to 900°C at a rate of 10°C/min.

X-ray Diffraction (XRD) measurements were performed using an XPert Pro-Philips diffractometer. The analysis was conducted over a 2Θ range of 10° to 80°, with a step size of 0.02° 2Θ and a time per step of 1 second.

### 2.3. Cloning of the VEGFC Expression Construct

The hygromycin resistance gene form pCoHygro (Invitrogen/ThermoFisher) was subcloned as a SapI/AccI fragment into pMT/BiP/V5-His C (Invitrogen/ThermoFisher), resulting in plasmid pMT-BiP-Hygro. To change the frame of the BglII restriction site with respect to the BiP signal peptide, we cloned annealed primers 5’-TGGCCTTTGTTGGCCTCTCGCTCGGA-3’ (lab-internal number M18800) and 5’-GATCTCCGAGCGAGAGGCCAACAAAGGCCACGA-3’ (lab-internal number M18801) into SfiI/BglII-opened pMT-BiP-Hygro. The resulting plasmid was named pMT-Ex and contains the BiP signal peptide, an extensive multiple cloning site followed by sequences for an optional V5 and hexahistidine tag, and is available via Addgene (#231106). The VEGF-C cDNA was finally inserted as a BamHI/ScaI fragment from pFB1-melSP-dNdC-C1HIS-hVEGF-C,^37^ into BglII/PmeI-opened pMT-Ex. The full sequence of the expression vector is available from Genbank (pMT-Ex-dNdC-hVEGF-C-H_6_, accession number PQ641246).

### 2.4. VEGFC Production and Purification

VEGFC was produced in Drosophila S2 cells stably transfected with expression vector pMT-Ex-dNdC-hVEGF-C-H_6_ (Genbank PQ641246) and purified as described by Karpanen et al.^38^ with the following modifications: Cells were grown in suspension culture in HyClone SFX-Insect cell culture medium (Cytiva, SH30350.02) under hygromycin selection (300 µg/ml) and induced with 1 mM CuSO_4_ at a density of 8 Mio. cells/ml. After 4 days of induction, the conditioned cell culture medium was separated from the cells by centrifugation for 5 minutes at 250 g. Imidazol was added to a final concentration of 10 mM, and the pH was adjusted to 7.4 with diluted NaOH. The supernatant was cleared by centrifugation (13000 g for 20 min). Excel sepharose (Cytiva) was added to the supernatant, and batch binding occurred under gentle agitation overnight at +4°C. After settling, the resin was recovered and applied to a 10-mm Diba Omnifit EZ column (Cole-Parmer). The column was washed with 20 mM imidazole, and elution was performed with 500 mM imidazole. The peak fraction was concentrated by ultrafiltration to 5 ml (Amicon Ultra Centrifugal Filter, 10 kDa MWCO, Millipore) and resolved on a size exclusion chromatography column (HiLoad 26/600 Superdex 200 pg, Cytiva) using PBS as buffer. The dimeric VEGFC peak (see Figure S1) was collected and sterilized via a sterile 4-mm PVDF syringe filter (Millex-GV, 0.22 µm, Millipore), and the concentration determined as 1.6 and 1.8 mg/ml by UV absorbance at 280 nm as measured by a DS-11 spectrophotometer (DeNovix) for the two gel filtration runs, respectively.

### 2.5. LNPs Synthesis

For nanoparticle synthesis, 15 mg of extracted lignin powder was mixed with 3 ml of acetone. After vortexing for 30 minutes to enhance the solubility of the lignin, the mixture was passed through a 0.45 µm polytetrafluoroethylene (PTFE) filter, and 1 ml of the filtrate was placed in a 15 ml polypropylene tube. Weighing the dry filter before and after filtration confirmed that there was only a negligible loss of lignin. Subsequently, 9 ml of water was added drop by drop at a rate of 11 ml/min under constant stirring. Finally, the suspension with a concentration of approximately 500 µg/ml was set aside for 1 hour before characterization.^39^

### 2.6. LNPs Characterization

The determination of size, zeta potential, and polydispersity index (PDI) was carried out using Dynamic Light Scattering (DLS) with a Zetasizer Nano ZSP (Malvern Instruments Ltd). The instrument was fitted with a 633 nm He−Ne laser, and measurements were performed at a backscattering angle of 173° using a clear disposable zeta cell cuvette. For the measurements, a working solution of 4 µg/ml was prepared from the 500 µg/ml stock solution.

To examine the morphology of the synthesized nanoparticles, both scanning electron microscopy (SEM) and transmission electron microscopy (TEM) were employed. SEM analysis was performed using a field-emission gun scanning electron microscope (FEG-SEM, Cambridge Leo Supra 35, Carl Zeiss). For TEM analysis, 5 µL of sample was deposited and dried on a Formvar carbon-coated copper grid (FCF-400-Cu, Electron Microscopy Sciences) and imaged on a Hitachi HT7800 Transmission Electron Microscope at 100 kV acceleration voltage.

### 2.7. Loading VEGFC Protein on LNPs

Drugs can be passively loaded onto nanoparticles by adding them to a solution containing previously prepared nanoparticles or by adding them to the reaction mixture during assembly.^40^

After examining different ratios, a nanoparticle-to-VEGFC ratio of 4:1 was found to give the best loading results, and two distinct loading approaches were employed.

The post-synthesis loading (PSL) approach, based on the work of Figueiredo et al. with some modifications, involved overnight shaking of a solution containing 30 µg/ml VEGFC and 120 µg/ml nanoparticles.^17^ Specifically, 3.75 µl of VEGFC from the 1.6 mg/ml stock solution was mixed with 48 µl of LNPs from a 500 µg/ml stock solution. The volume was adjusted to 200 µl with sterilized MQ water and maintained in a rotary shaker for 24 hours (180 rpm, 1.9 or 2.5 cm throw, Figure 1A).

**Figure 1.**
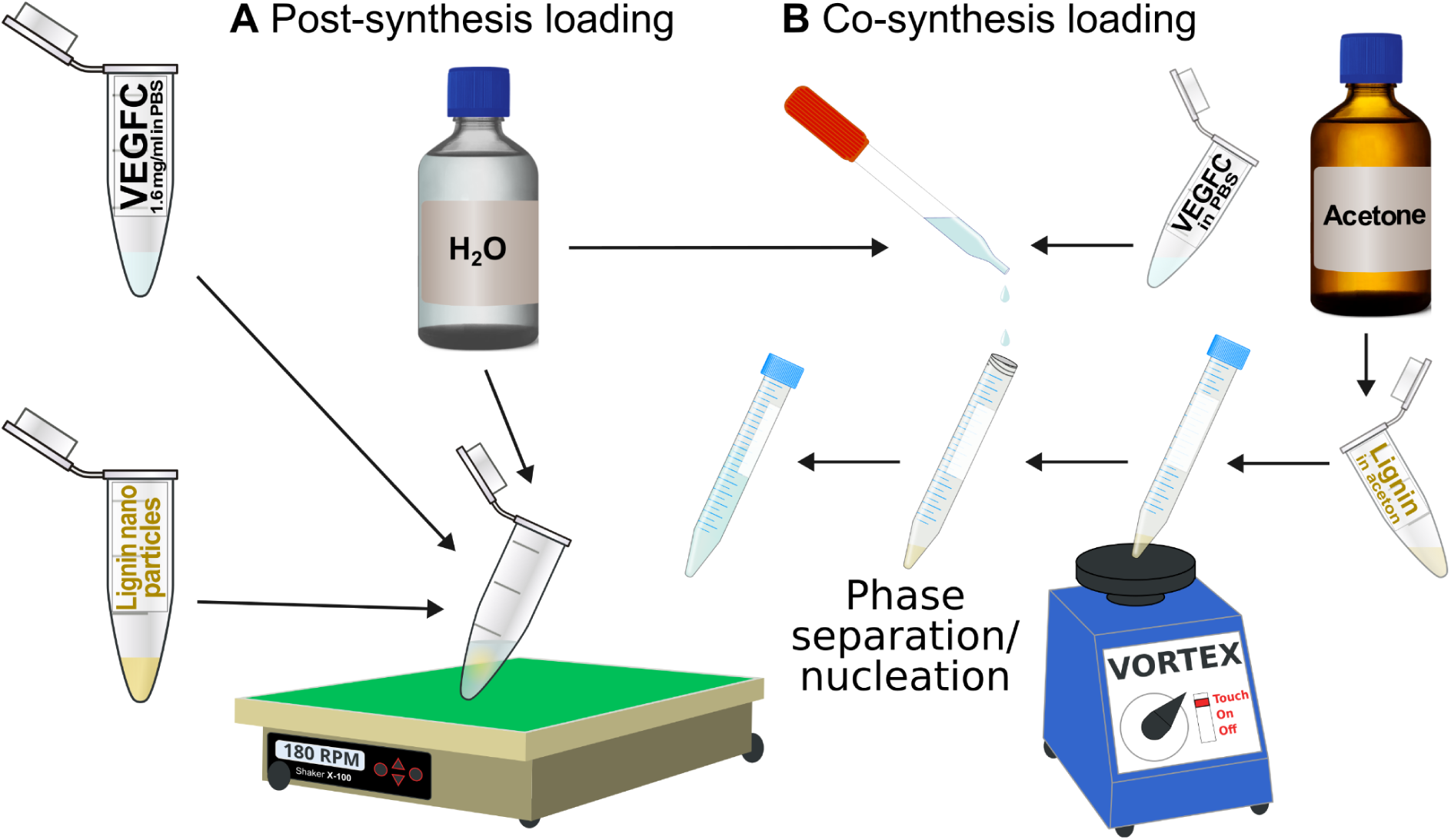
(**A**) Post-synthesis loading was done by coincubating LNPs with VEGFC overnight in 200 µl at 25°C under agitation (30 µg/ml VEGFC, 120 µg/ml nanoparticles). (**B**) Co-synthesis loading was achieved by inducing nucleation by reducing the solubility of a lignin/VEGFC mix by adding water.

The co-synthesis loading (CSL) approach, adapted from Zhou et al., involved incorporating VEGFC during nanoparticle synthesis.^41^ To achieve this, 15 mg of lignin powder was dissolved in 3 ml of acetone to achieve a concentration of 5 mg/ml. The mixture was vortexed for 30 minutes and filtered using a 0.45 µm PTFE syringe filter. Then, 240 µl of the filtered lignin was placed into a polypropylene tube. A mixture of protein and water was prepared by combining 166.7 µl of VEGFC (from the 1800 µg/ml stock solution) with 9.6 ml of water. This mixture was gradually added dropwise to the filtered lignin immediately after homogenously dispersing the suspension. As the lignin separated, nanoparticles formed, resulting in about 10 ml suspension with a final concentration of 120 µg/ml nanoparticles and 30 µg/ml VEGFC (Figure 1B).

100 µL of the final product from each approach was centrifuged at 16100g for 5 minutes. Following centrifugation, 80 µL of the supernatant was removed for the analysis of free VEGFC using reverse phase high-performance liquid chromatography (HPLC; Agilent 1100 series, Agilent Technologies, USA).^17^ After the supernatant was removed for HPLC analysis, the precipitate was resuspended in water containing 2% acetone to induce partial detaturation and release of the attached VEGFC. The concentration of VEGFC in the precipitate was then measured by HPLC. Knowing the concentrations of free VEGFC, attached VEGFC, and total VEGFC, the loading capacity of the nanoparticles, as well as the yield, were calculated.

HPLC was conducted to detect and quantify VEGFC using a mobile phase of 35% trifluoroacetic acid (0.1%) and 65% acetonitrile. A 15 µL sample was injected, and separation was achieved with a 75 × 4.6 mm column (Kinetex 2.6 µm XB-C18 100Å). A calibration curve for VEGFC was established, covering the range from 10 to 100 µg, and chromatograms were obtained for VEGFC, LNPs, and VEGFC-loaded nanoparticles for peak identification. The calibration curve was used to quantify the free VEGFC (Figure 2).

**Figure 2.**
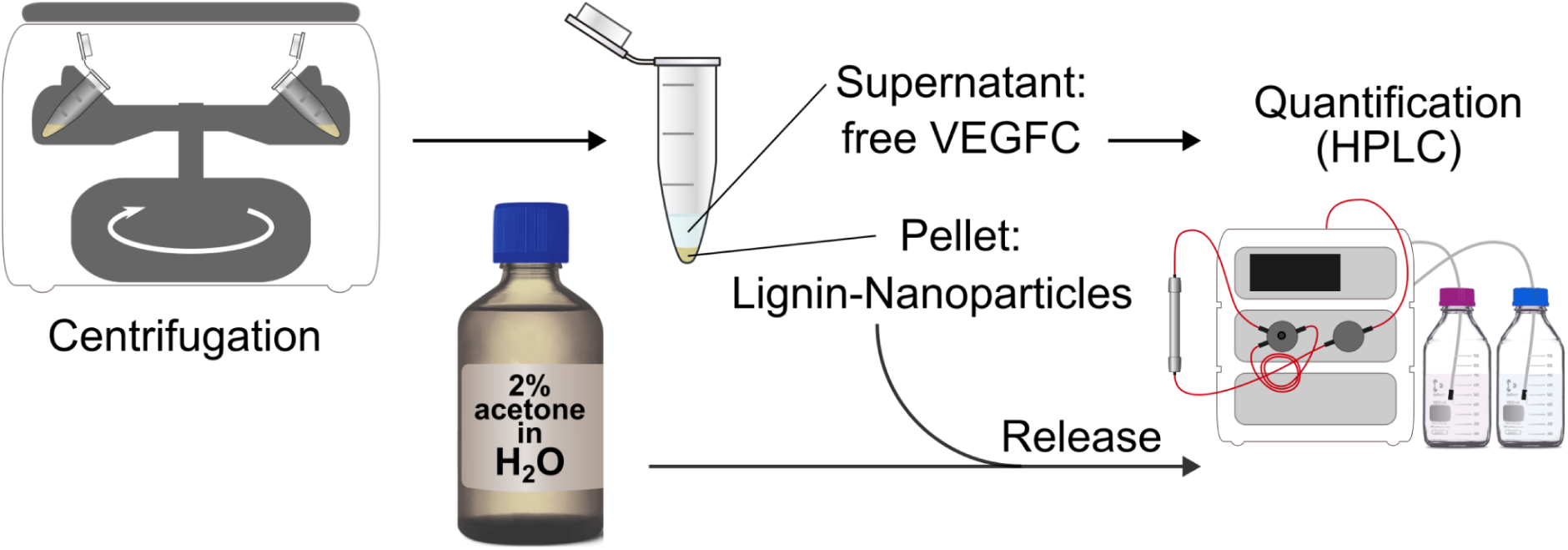
Loading was characterized by isolating the nanoparticles by centrifugation and quantifying free VEGFC from the supernatant and bound VEGFC from the pellet after releasing it with 2% acetone in water. The amount of attributed loss or degradation of VEGFC is then calculated by subtracting the sum of free VEGFC and bound VEGFC from the total VEGFC.

### 2.8. Growth Factor Release Assay

Co-synthesis-loaded LNPs appeared to be superior to post-loaded LNPs, and therefore, we used for the growth factor release assay co-synthesis-loaded LNPs. The release profile was assessed in two modes: one where free VEGFC was not removed, and another where free VEGFC was removed prior to the release.

The release profile of VEGFC from the nanoparticles was examined in two release media: HBSS-HEPES (Hank’s Balanced Salt Solution with HEPES (4-(2-hydroxyethyl)-1-piperazineethanesulfonic acid, pH 7.4); and HBSS-MES-T (Hank’s Balanced Salt Solution with MES (2-(N-morpholino)ethanesulfonic acid, pH 5.5, 2% Tween 80).

For the suspension without removing free VEGFC, 1 ml of a nanoparticle suspension containing 120 µg/ml of nanoparticles and 30 µg/ml of VEGFC were mixed with 2 ml of the release medium, resulting in a final VEGFC concentration of 10 µg/ml. To remove free VEGFC, the nanoparticle suspension containing 120 µg/ml of nanoparticle and 30 µg/ml of VEGFC was centrifuged at 13,200 rpm for 5 minutes. The VEGFC concentration of the supernatant was quantified using HPLC, and the concentration of the attached VEGFC was calculated. To create the control sample containing an equal amount of free VEGFC, 50 µl of VEGFC from the 1600 µg/ml stock solution were diluted with release medium into a final volume of 8 ml.

To evaluate the release profile of VEGFC from the LNPs, a series of timed withdrawals were performed. Specifically, 100 µl samples were taken from both the experimental and control groups at 0 minutes, 5 minutes, 15 minutes, 30 minutes, 2 hours, 4 hours, 6 hours, and 24 hours after exposure to the release medium. The release media were kept on a stirrer plate at 37°C and stirred at 170 rpm. Following each withdrawal, an equal volume of fresh, preheated release medium (100 µl) was added to maintain an equal volume (Figure S2).

The collected samples underwent centrifugation at 16100 g for 5 minutes to separate the supernatant from the nanoparticles. Subsequently, 80 µl of the supernatant was transferred into HPLC well plates for the quantification of free VEGFC. For quantifying each sample in both media, calibration curves for HBSS-HEPES and HBSS-MES-T were established.^17^

### 2.9. Nanoparticle Stability in Blood Plasma

The stability of nanoparticles in human blood plasma at 37°C was assessed by measuring changes in size, PDI, and zeta potential over 90 minutes. Plasma from anonymous donors, obtained with ethical approval, was cleared by centrifugation for 4 minutes at 25°C and 1915 g. LNP solution was mixed with plasma to a concentration of 40 µg/ml and incubated at 37°C. Aliquots were taken at specified intervals and diluted for Dynamic Light Scattering (DLS) analysis. A control with plasma alone was also tested, with all experiments performed in triplicates.^17^

### 2.10. Cell Proliferation/Viability Assessment

To assay the biological activity of VEGFC released from LNPs, the VEGFC-sensitive cell line Ba/F3-VEGFR-3/EpoR was used.^42^ Cells were cultured in Dulbecco’s Modified Eagle Medium (D-MEM) supplemented with 10% fetal calf serum (FCS), interleukin-3 (IL-3, 2 ng/ml), VEGFC (50 ng/ml), and zeocin (200 µg/ml). After harvesting and washing, cells were resuspended in D-MEM with 10% FCS and seeded at 5,000 cells/well in 96-well plates. Cells were treated with VEGFC-loaded nanoparticles after the removal of excess free VEGFC, nanoparticles alone, VEGFC, or control medium, and incubated at 37°C with 5% CO₂ for 10 days. Metabolic activity was assessed daily, similar to the previously published protocol,^42^ using the chromogenic substrate MTT followed by cell lysis and measurement of absorbance at 570 nm using a microplate reader to determine metabolic activity as a combined proxy for proliferation and viable cell count.

### 2.11. VEGF-C Stability Assays

293F cells that stably express pro-VEGF-C (a kind gift from Sawan K. Jha) were grown in 6-well plates in D-MEM 10% FCS. To estimate the stability of VEGFC against proteolytic digestion, the culture volume was reduced to 1.5 ml/well, and we exchanged 50% of the medium with methionine- and cysteine-free D-MEM supplemented with 10 µCi EasyTag Express [35S] protein labeling mix (PerkinElmer/Revvity) per ml. Thermolysin, trypsin, and proteinase K (Sigma-Aldrich/Merck, T7902, T4799, and P6556, respectively) were added to final concentrations of 0, 1, 5, 30, and 150 ng/ml. After 3 days of culture, supernatants were removed and centrifuged twice at 400 and 11000 g, respectively. 2 µg of VEGFR-3/Fc,^43^ 0.5 mg BSA, and 45 µl of a 50% Protein A sepharose CL-4B (PAS, GE Healthcare/Cytiva) slurry in PBS were added to each sample and incubated at 4°C with gentle agitation overnight. The PAS was separated by centrifugation at 1500g for 5 minutes and washed three times with PBS. 25 µl of 2x reducing Laemmli Standard Buffer were added, and after heating for 5 minutes at 95°C, samples were resolved on a 4-20% PAGE gel (Mini-PROTEAN TGX, Bio-Rad). The gel was dried, exposed to a Fujifilm phosphoimager plate overnight, and read with a Typhoon FLA 7000 scanner (GE Healthcare/Cytiva).

To evaluate the stability of VEGFC against prolonged storage, purified mature VEGFC was stored at 4, 21, 37, or 50°C for 0, 1, 2, 4, and 8 days. To test the resilience of VEGFC against freezing-thawing, 5 µl of purified mature VEGFC in 1.5-ml Eppendorf tubes was cycled between room temperature and dry ice (5 minutes each). The samples were used to support cell growth and survival of Ba/F3-VEGFR-3/EpoR cells as described above, except that the cell seeding density was 20000 cells/well and that a single assay was performed 48 h after seeding (Figure S3).

### 2.12. Statistical Analyses

Statistical analysis was done in Gnumeric 1.12.56. Error bars in the figures denote the standard deviation (STDEV) except for the release assays, which show the standard error of the mean (SEM).

## 3. Results

### 3.1. Lignin Characterization Confirms Its Structural Identity

The amount of lignin obtained was 0.37 g from an initial 4 g wood sample, with a yield of 45.7% calculated according to Jimenez et al. (2007).^6^ The FTIR spectrum of the extracted lignin is depicted in Figure S4A, displaying characteristic absorption bands indicative of lignin’s functional groups. The UV-Vis absorption spectrum of lignin is shown in Figure S4B and features the prominent absorption peak around 280 nm.^44,45^ Simultaneous Thermogravimetric Analysis and Differential Scanning Calorimetry (TG-DSC) measurements were performed in air to study the thermal stability of the samples (Figure S5). The XRD pattern of lignin provides insights into its crystalline and amorphous structures. The XRD pattern shows a broad peak centered around 22° 2θ ( Figure S6).

### 3.2. Size, Surface Charge, and Morphology Confirm LNP Properties

After synthesizing the LNP, the size and zeta potential were determined by dynamic light scattering. The results were 142±62 nm and -40±8 mV for the Z-average and the zeta potential, respectively ( Figure S7). The PDI was 0.126, suggesting a relatively narrow size distribution and indicating a fairly monodisperse sample.^46^ The SEM image of LNPs shows an average diameter of 194±6. SEM images revealed the spherical morphology and rough surface texture of the LNPs. The uniformity in the TEM images supports the low PDI value of 0.126 from the DLS analysis, indicating a narrow size distribution (Figures 3A and 3B).

**Figure 3.**
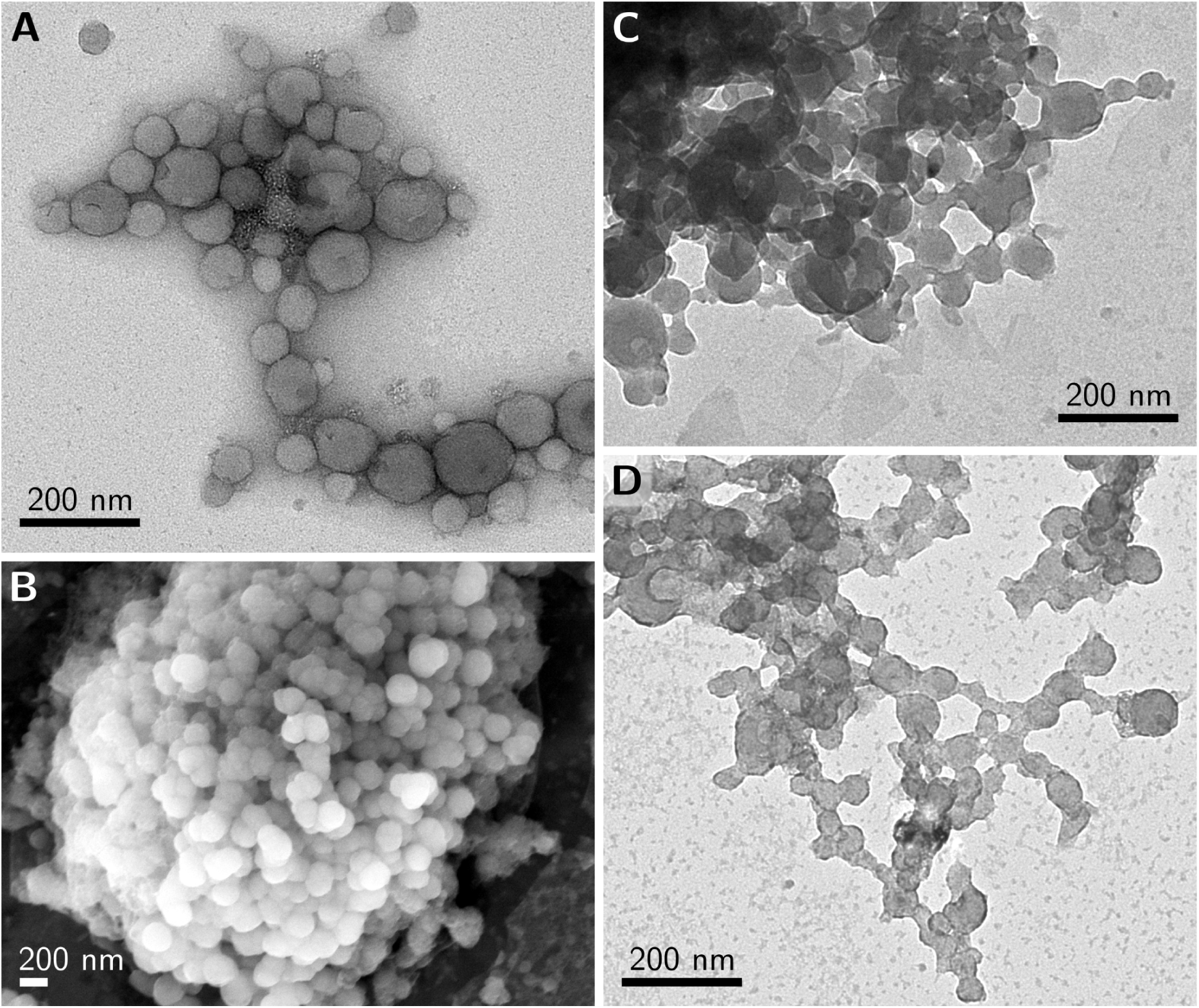
Morphological characterization of LNPs and VEGFC-loaded LNPs using electron microscopy. **(A)** TEM image of LNPs showing their spherical structure and aggregation tendency **(B)** SEM image of LNPs, highlighting their surface topology and size distribution. **(C)** TEM image of VEGFC-loaded LNPs prepared via post-synthesis loading, displaying particle clustering and aggregation **(D)** TEM image of VEGFC-loaded LNPs using co-synthesis loading, showing a more interconnected and network-like nanoparticle arrangement (For the size distribution of LNPs, see also Figure S7).

### 3.3. TEM Characterization of VEGFC-Loaded LNPs Reveals Well-Defined and Better-Dispersed Nanoparticles in CSL Approach

The loading capacities of LNPs using post- and co-synthesis approaches were determined. The results, including the concentrations of free VEGFC and bound VEGFC for each approach, are summarized in Table 1.

**Table 1.** Capacities of the different loading approaches. Detected free VEGFC was subtracted from the initial concentration (30 µg/ml, or 3 µg mass in 100 µl, as used for analysis), and the amount of VEGFC incorporated into the lignin nanoparticles was determined.

| Approach | Post-synthesis loading (PSL) | Co-synthesis loading (CSL) |
| --- | --- | --- |
| Conc. of free VEGFC | 9.02 $\mu\text{g/ml}$ (0.902 $\mu\text{g}$) | 2.28 $\mu\text{g/ml}$ (0.228 $\mu\text{g}$) |
| Conc of bound VEGFC | 8.92 $\mu\text{g/ml}$ (0.892 $\mu\text{g}$) after dissolving the precipitate in 2% acetone water | 22.5 $\mu\text{g/ml}$ (2.25 $\mu\text{g}$) after dissolving the precipitate in 2% acetone water |
| Loading percentage | $8.92/30 \times 100 = 29.73\%$ | $22.5/30 \times 100 = 75\%$ |

The TEM images for PSL show VEGFC-loaded LNPs with some aggregation, as well as a broader size distribution (Figure 3C). In contrast, the TEM images for CSL depict nanoparticles that appear more distinguishable, with a more defined spherical shape and better dispersion (Figure 3D). While these images suggest differences in morphology, dispersion, and structural integrity after loading, they do not directly indicate variations in VEGFC loading.

### 3.4. Slightly Enhanced Release at pH 5.5 Compared to pH 7.4

The release profiles of VEGFC from the LNPs, which were loaded using the CSL approach, were analyzed under physiological (pH 7.4) and endosomal (pH 5.5) conditions (Figure 4).

**Figure 4.**
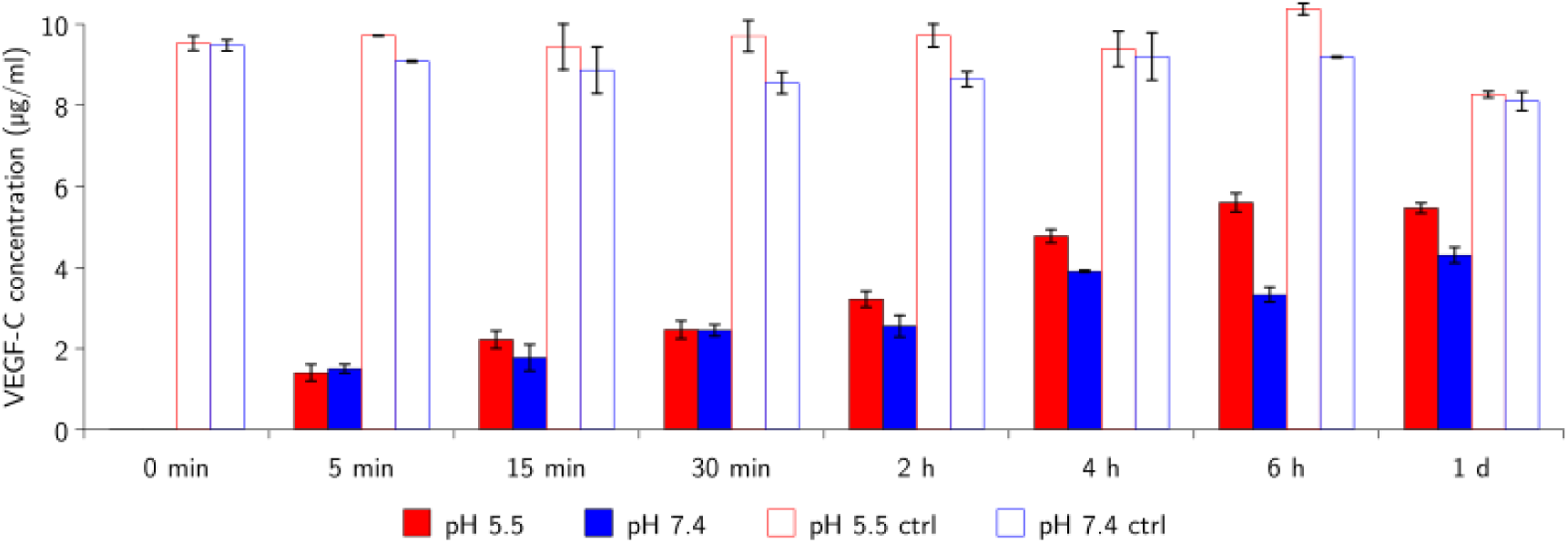
Release assays for VEGFC. The release profiles of VEGFC from the co-synthesis loaded LNP were analyzed under physiological (pH 7.4) and endosomal (pH 5.5) conditions (also see Figure S2).

The attached amount of nanoparticle-attached VEGFC was quantified as 24.5 µg/ml by subtracting the HPLC-quantified free VEGF-C (5.5 µg/ml) from the original sample concentration of 30 µg/ml. After diluting into the release medium, the final concentration of the nanoparticle-attached VEGFC in the release media was 8.16 µg/ml.

The sample, after removing the free VEGF in pH 7.4, demonstrated a steady increase from 0 µg/ml to 2.5 µg/ml by 2 hours and peaking at around 4.30 µg/ml after 24 hours, indicating a controlled release. Additionally, a mild decline from 4 to 6 hours was noted at pH 7.4.

A similar release pattern was observed at the endosomal pH of 5.5, but with a higher and faster release. Release in pH 5.5, started at 0 µg/ml and showed a controlled, gradual release, reaching approximately 2.23 µg/ml by 15 minutes. This sample continued to exhibit a steady release up to 24 hours without significant decline and peaked at 5.63 after 6 hours, indicating a sustained release pattern.

From another perspective, at pH 7.4, an initial attached VEGFC concentration of 8.16 µg/ml results in the release of 4.30 µg/ml after 24 hours, indicating a release percentage of approximately 53%. Similarly, at pH 5.5, 5.60 µg/ml is released from the same initial concentration over 24 hours, indicating a release percentage of approximately 69%.

The control average shows a relatively stable concentration of VEGFC over time, starting around 10 µg/ml and maintaining this level with minor fluctuations. A slight decline was observed, likely indicating the absence of nanoparticle protection, which might lead to the gradual degradation or loss of VEGFC over time.

### 3.5. In Vitro Assay

#### 3.5.1. Potential Interactions Between LNPs and Plasma Proteins Reflected in Size Distribution

The stability of the loaded nanoparticles in plasma was assessed to evaluate their potential for systemic circulation. This analysis is crucial for understanding the degradation or aggregation behavior of nanoparticles in a biological matrix, as shown in Figure 5.

**Figure 5.**
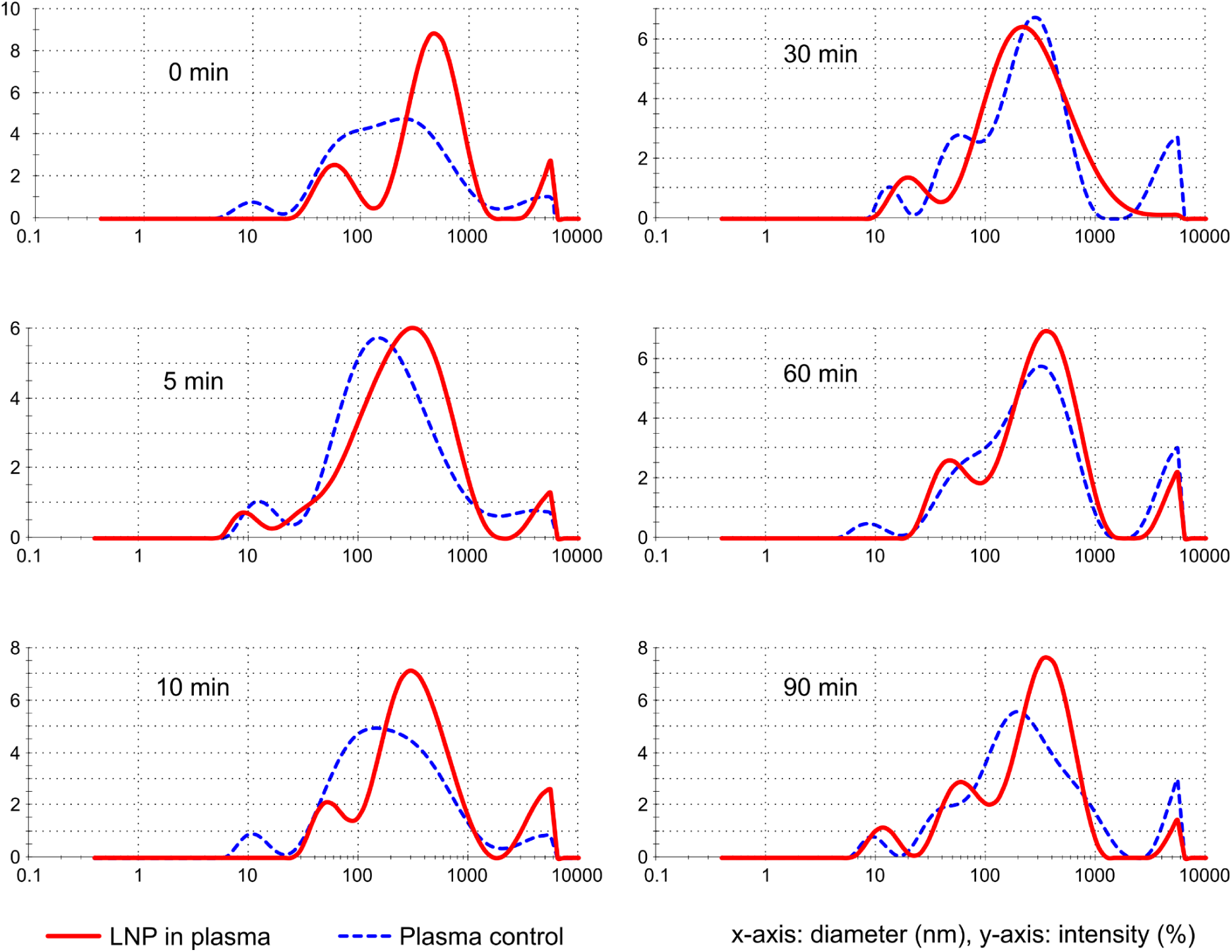
Changes in the size of LNPs in plasma over time. Demonstrating their dynamic behavior and moderate stability under physiological conditions, with plasma control size changes used as a baseline to compare the stability and behavior of LNPs in plasma.

The stability of LNP in plasma was evaluated using DLS to monitor changes in size distribution and zeta potential over time (Table T1). These assessments were conducted to understand how LNPs interact with plasma components, which is crucial for their potential application as drug delivery systems.

#### 3.5.2. Possible Sustained Release and Enhanced Bioavailability of VEGFC with LNP

The VEGFC release from the loaded nanoparticles was evaluated using cell viability assays. The Ba/F3 assay measures proliferation and viability using cytokine-dependent Ba/F3 cells. The readout OD values reflect metabolic activity, integrating both proliferation/survival and viability. The different response patterns of cells after exposure to VEGFC alone and VEGFC-loaded LNP were studied. Absorbance measurements as a proxy for cell survival and proliferation were recorded over 10 days for different VEGFC concentrations of three samples: VEGFC alone (positive control), LNP + VEGFC, and LNP alone (negative control) (Figures 6 and 7).

**Figure 6.**
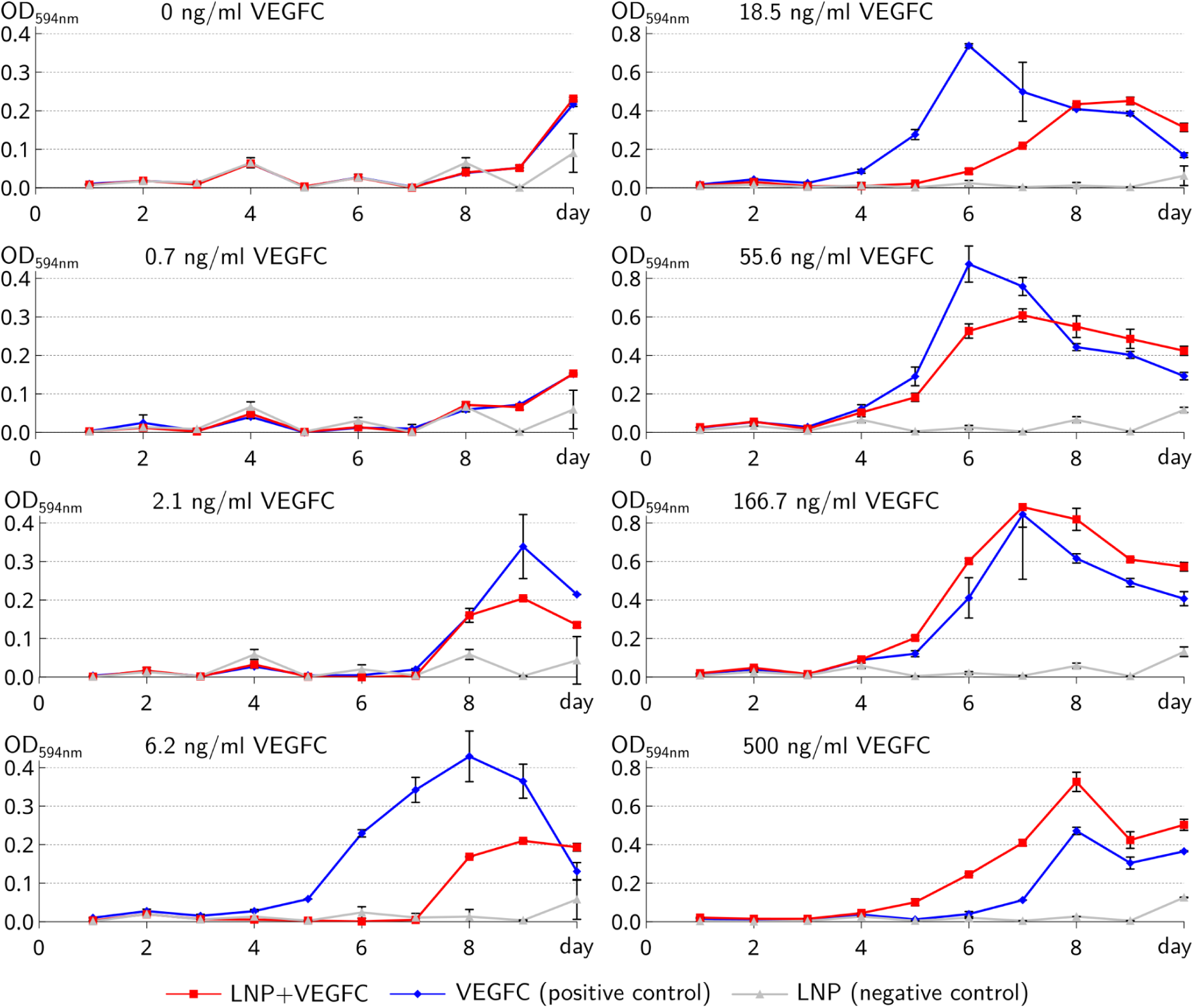
Proliferation/viability of VEGFC-responsive Ba/F3 cells over a period of 10 days. Absorbance was measured daily for VEGFC alone (positive control, shown in blue), LNP + VEGFC (shown in red), and LNP alone (negative control, shown in gray). At intermediate concentrations, the LNP + VEGFC condition exhibited a delayed but sustained increase in absorbance, suggesting controlled VEGFC release compared to the rapid peak observed in the VEGFC positive control.

**Figure 7.**
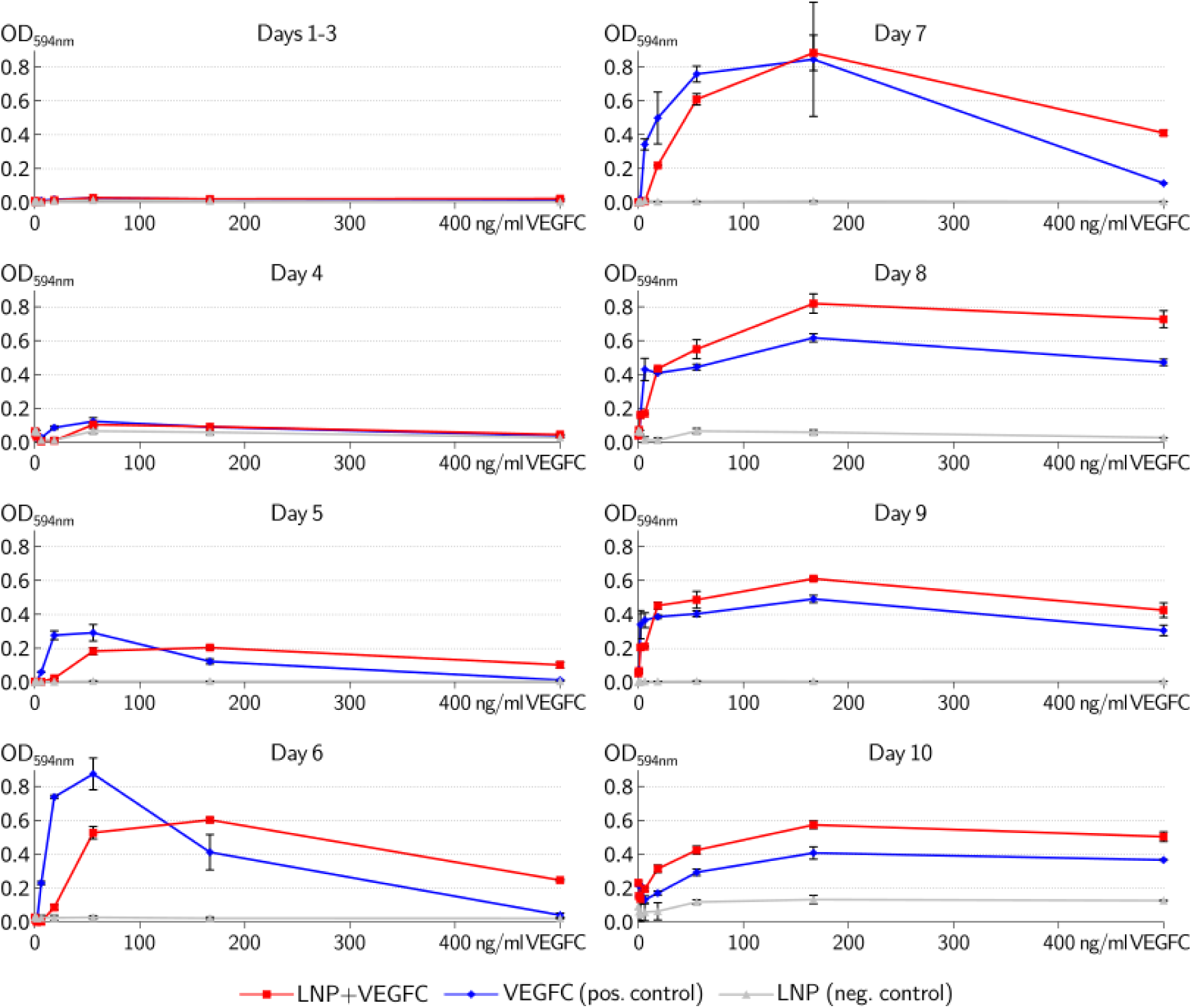
Proliferation/viability of VEGFC-responsive Ba/F3 cells in response to different VEGFC concentrations. From day 4 onward, LNP + VEGFC demonstrated, at intermediate and higher concentrations, a more sustained increase in cell proliferation compared to the VEGFC positive control, suggesting prolonged VEGFC release from nanoparticles. VEGFC concentrations above 200 ng/ml seem to have a negative effect on cell survival/proliferation, which appears less pronounced for the LNP + VEGFC samples.

The response increased with VEGFC concentration, reaching its maximum around 100 ng/ml. At very low concentrations (0–2.1 ng/ml), minimal to moderate increases were observed, with no clear differences between VEGFC and LNP + VEGFC. At mid-range concentrations (6.2–18.5 ng/ml), peak absorbance occurred earlier in the VEGFC control (day 6–8), while LNP + VEGFC exhibited delayed but sustained peaks. At higher concentrations (55.6–500 ng/ml), both conditions showed strong absorbance increases, with LNP + VEGFC peaking slightly later but maintaining a higher level at the highest concentration of 500 ng/ml. Across all tested concentrations, the LNP + VEGFC condition consistently exhibited a delayed peak and a more gradual decline compared to the VEGFC control.

From day 1 to 3, absorbance values are low and nearly identical across all conditions, indicating minimal changes. From day 4 onward, absorbance increases in both the VEGFC positive control and Lignin NP + VEGFC, with distinct differences observed between VEGFC alone and VEGFC-loaded nanoparticles.

## 4. Discussion

Therapeutic growth factor delivery for vascular regeneration has been challenging. Although the feasibility of generating new vessels (both blood and lymphatic) has been experimentally demonstrated,^47^ none of the clinical trials has provided convincing evidence of a significant therapeutic benefit.^48^ A possible explanation has been the technical inability to deliver the growth factors in the right amount to the right location over a sustained period.^49^ While single injections of VEGFC have shown beneficial effects in some models of heart regeneration after myocardial infarction,^50^ sustained-release systems are likely to provide better results. Using lymph node transplantation to jump-start the regeneration of lymphatic drainage in breast cancer-associated lymphedema is perhaps successful due to the lymph nodes or recruited macrophages providing VEGF-C,^51^ and overprovisioning VEGF-C could increase the success rate of this intervention.^52,53^ In this study, we wanted to elucidate whether lignin nanoparticles could provide a sustained source of VEGFC to aid in vascular regeneration.

DLS analysis revealed that the primary peak of LNP was centered at 165 nm with 100% intensity, confirming that most nanoparticles are around this size, with a standard deviation of 65 nm indicating some variation but maintaining a consistent size range. The Zeta Potential analysis showed a mean value of -40±8 mV, reflecting the stability of the LNP dispersion. A high magnitude of zeta potential indicates good stability due to electrostatic repulsion between particles, preventing aggregation. These findings show that the LNPs have a small and uniform size distribution with good colloidal stability. The low PDI value supports the monodispersity of the sample, and the high negative zeta potential suggests stable dispersion, similar to the findings of Ortega-Sanhueza et al. (2024), who reported lignin nanoparticles with sizes ranging from 113 to 154 nm, a PDI of 0.09 to 0.26, and a zeta potential of -23 to -41 mV, indicating stability and resistance to agglomeration.^54^

SEM also indicates that the nanoparticles are well-dispersed and stable, suitable for applications requiring consistent particle sizes. SEM images revealed the spherical morphology and rough surface texture of the LNPs, indicating a high surface area beneficial for drug loading applications. However, the significant aggregation observed in SEM is likely due to the drying process required for analysis, which does not accurately reflect their true dispersion state in a liquid medium.

In contrast, TEM images provided clearer insights into individual nanoparticles, displaying well-defined, uniformly spherical particles and better dispersion. The particles are observed to be in the range of 150-200 nm. The morphology of the nanoparticles observed in our study appears consistent with previous findings.^55^

Drugs can be loaded onto nanoparticles by adding them after nanoparticle synthesis or by incorporating them during the synthesis reaction.^40^ In the current study, the loading efficiency of VEGFC onto LNPs was evaluated using both approaches. PSL resulted in a loading efficiency of almost 30%. In contrast, CSL achieved a loading efficiency of 75%. The lower efficiency observed with PSL in this approach could be due to several factors. Primarily, the interaction between VEGFC molecules and the nanoparticle surface might be limited after the nanoparticles have already formed. This can lead to less effective encapsulation and potential loss of VEGFC during the loading process. This method does not allow VEGFC to integrate fully into the nanoparticle matrix, thus reducing its loading capacity.^56^ In addition to more effectively trapping VEGFC molecules within the nanoparticles, CSL reduced the likelihood of VEGFC loss and enhanced both the stability and binding efficiency within the nanoparticles. We speculate that the phase separation process during CSL further enhanced the trapping of VEGFC molecules within the forming nanoparticles.^39^ Integrating protein cargo during nanoparticle synthesis generally offers higher encapsulation efficiency, stability, and potentially better therapeutic outcomes.^57^ The particle surface charge/zeta potential can significantly impact loading efficiencies.^58^ Similar to most other VEGF family members, the VEGFC form we used carries a slightly positive charge at physiological pH,^59,60^ aiding in the interaction with the negative surface charge of the LNPs. However, we did not see a significant delayed release in pH 5.5 versus 7.4. TEM images from PSL reveal VEGFC-loaded LNPs with aggregation and broader size variations. On the other hand, CSL shows more spherical and well-dispersed nanoparticles. This improvement could be due to the simultaneous synthesis and loading of VEGFC, allowing the nanoparticles to form around the VEGFC molecules more effectively. The TEM images of CSL show similarities with the findings in the articles by Valo et al. (2010) and Valo et al. (2013).^61,62^ The optimized sample, CSL, was chosen for the release assay. It appears that this sample exhibited a controlled and sustained release due to gradual diffusion from within the nanoparticles. The pH-dependent release behavior observed in HBSS-HEPES (pH 7.4) and HBSS-MES (pH 5.5) demonstrated faster release at acidic pH, simulating the endosomal microenvironment. This pH-responsive release is crucial for targeted drug delivery, as shown by Figueiredo et al. (2017) and Kumari et al. (2010).^63^

The Z-average size of LNPs in plasma initially increased from 141.2 nm to 213.3 nm, indicating nanoparticle presence, and decreased over time to 156.0 nm at 90 minutes as they stabilized, suggesting dynamic interactions with plasma proteins. Secondary peaks in the nanoparticle samples, more intense than in plasma alone, indicate the formation of nanoparticle-protein complexes, with peaks at 61 nm and 58 nm. The consistent zeta potential of around -20 mV for both plasma alone and nanoparticle-plasma mixtures supports the moderate stability of these complexes, showing that although nanoparticles actively interact with plasma proteins, this does not significantly alter the system’s overall surface charge. In the bloodstream, nanoparticles interact with proteins, forming a coating (corona) that influences how they interact with tissues and affects their overall behavior and fate.^64^ The protein corona changes the nanoparticle properties, influencing their potential use in biomedical applications.^65^ Moreover, it reduces their stealth and targeting abilities, leading to either prolonged or quicker clearance by the immune system.^66^ LNP stability in plasma, indicated by the gradual size difference and stable zeta potential, suggests partially stable nanoparticle-protein complexes. Size distribution changes and secondary peak intensity reflect active nanoparticle-protein interactions, crucial for drug delivery, and consistent with studies on protein coronas.^67,17^.

We modified the traditional Ba/F3 MTT assay, which typically starts with much higher cell densities, and assays only once at 48 hours or less.^43,68,69^ Using a much lower initial cell density, we were able to follow the cell proliferation over a period of 10 days. At mid to high concentrations (6.2 ng/ml and above), the Lignin NP + VEGFC demonstrates better-sustained release and bioavailability compared to the VEGFC positive control. The delayed and sustained release profile was particularly evident at 6.2, 18.5, and 55.6 ng/ml, indicating that the controlled release provided prolonged availability of VEGFC at physiologically relevant concentrations (at and above the K_D_ value of ∼135 pM for the ligand/receptor interaction). Although the nanoparticles still enhanced bioavailability at higher concentrations, the differences became less pronounced, likely due to receptor saturation. The lignin negative control (Lignin NP without VEGFC) consistently shows low absorbance values, similar to the baseline proliferation level, indicating no major cytotoxic effects of the nanoparticles themselves on Ba/F3 cells (Figure S8). This suggests that the LNPs are biocompatible. Both VEGFC + NP and VEGFC alone exhibit similar kinetic patterns at concentrations of 166.7 and 500 ng/ml, with VEGFC + LNP consistently showing a slightly higher proliferation/viability than VEGFC alone. With a K_D_ of 135 pM (or 5.7 ng/ml)for the VEGFC ligand,^70^ the VEGFR-3 becomes saturated at higher concentrations, leading to a plateau effect where additional VEGFC, whether free or delivered via nanoparticles, does not significantly increase cell proliferation.^71,72^ However, the nanoparticle system may prevent toxic concentrations of VEGFC by providing controlled release, which was observed at the highest concentration.^73,74^ While in-vivo data shows that too high VEGFC concentrations are detrimental to lymphatic vessel health,^75^ an alternative explanation could be impurities in the VEGFC preparation that start to interfere with cell proliferation/viability at the highest concentration. Furthermore, nanoparticles may protect VEGFC from degradation, enhancing its bioavailability and stability. However, it is unlikely that this mechanism is a major contributor because mature VEGFC is a rather stable molecule. We show that it is relatively resistant to proteolytic attacks, including longer storage at elevated temperatures and limited exposure to proteinases (Figure S9). When tested for stability at different temperatures, VEGFC exhibited maximal biological activity in the Ba/F3-VEGFR-3/EpoR assay even after 8 days of storage at 37°C (Figure S3A). Interestingly, it was relatively sensitive to freeze-thawing cycles (Figure S3B). Even though there were clear differences in the bioassay between adding VEGFC as free protein and VEGFC-loaded nanoparticles, these were less pronounced than we had expected. One possible explanation is that VEGFC could by itself already associate with extracellular matrix (ECM) or cell surfaces to form a slow-release depot.^76^ Pro-VEGFC is able to interact with a plethora of ECM and cell surface proteins,^76^ and we speculate that the mature form of VEGFC might maintain some of the heparan proteoglycan affinity of proVEGFC.

## 5. Conclusion

LNPs demonstrate potential as drug delivery systems due to their ability to load VEGFC and release it in a controlled manner, thereby enhancing its availability and promoting cell growth, as shown in cell survival assays. However, the differences between VEGFC-loaded nanoparticles and free VEGFC were less significant than anticipated, indicating that further in vivo studies are required to better understand their behavior in complex biological systems and assess their safety and effectiveness for medical use.

Additionally, exploring the potential of LNPs for targeted drug delivery in specific diseases, including cancer and chronic conditions, could lead to more effective and personalized treatments. Investigating the use of LNPs in combination therapies, where multiple drugs or therapeutic agents are delivered simultaneously, may also open new avenues for enhancing treatment outcomes.

## Supporting information

Full size uncropped images

## Contributions

AMZ performed most of the experiments, analyzed the data, and wrote the manuscript; EG and RL contributed to lignin extraction and characterization, and nanoparticle design and characterization; HB designed and performed the bioassays, and prepared samples for releasing and TEM parts; A. Caravella helped with Lignin extraction from biomass; CD assisted with SEM imaging; EDB contributed to X-ray measurements; FD assisted with DLS setup for size and zeta potential; SS performed and repeated the stability testing of VEGFC; A. Correia helped with DLS and contributed to loading, releasing, and plasma assay parts; KR produced and purified the VEGF-C protein; ZG helped with the early discussion and review; AK performed TEM imaging and edited related section in the manuscript; TL designed a new approach for release study, interpreted TEM images and helped with manuscript editing; JK supervised the protein production and purification and contributed to manuscript writing; MJ supervised the study and helped with the bio- and stability assays and the manuscript writing.

## Acknowledgments

We thank Sawan K. Jha (University of Stanford, CA) for providing the stable 293F cell line that expresses pro-VEGFC and Kenny Mattonet (Max Planck Institute for Heart and Lung Research, Germany) for performing the first experiments to test the stability of VEGFC. We also thank the Electron Microscopy Unit (EMBI) at the Institute of Biotechnology, supported by HiLIFE and Biocenter Finland, at the University of Helsinki.

## Supplementary Figures and Tables

**Supplementary Figure S1.**
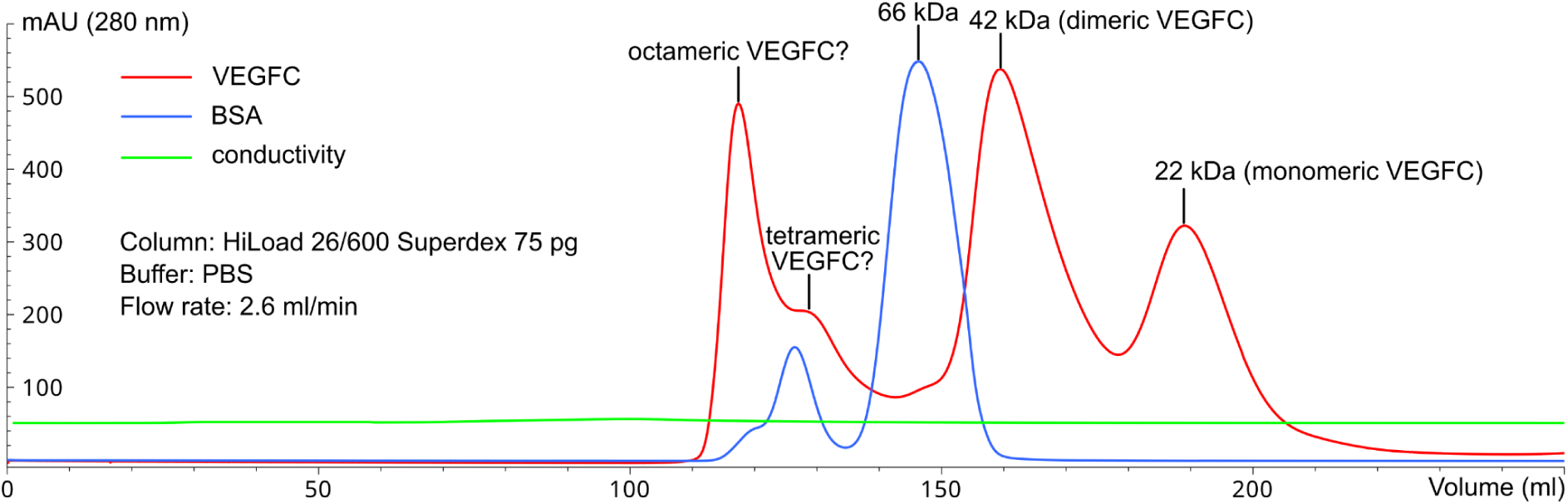
Size exclusion chromatography of mature VEGFC. VEGFC purified by IMAC was subjected to size exclusion chromatography. The elution profile of VEGFC indicated that insect cells produce an approximately equimolar mix of dimeric and monomeric species. Peaks were identified using BSA as a molecular weight marker. However, there is considerable batch-to-batch variation concerning the ratio between dimeric and monomeric VEGFC. We also observed higher molecular weight species that putatively corresponded to tetrameric and octameric assemblies.

**Supplementary Figure S2.**
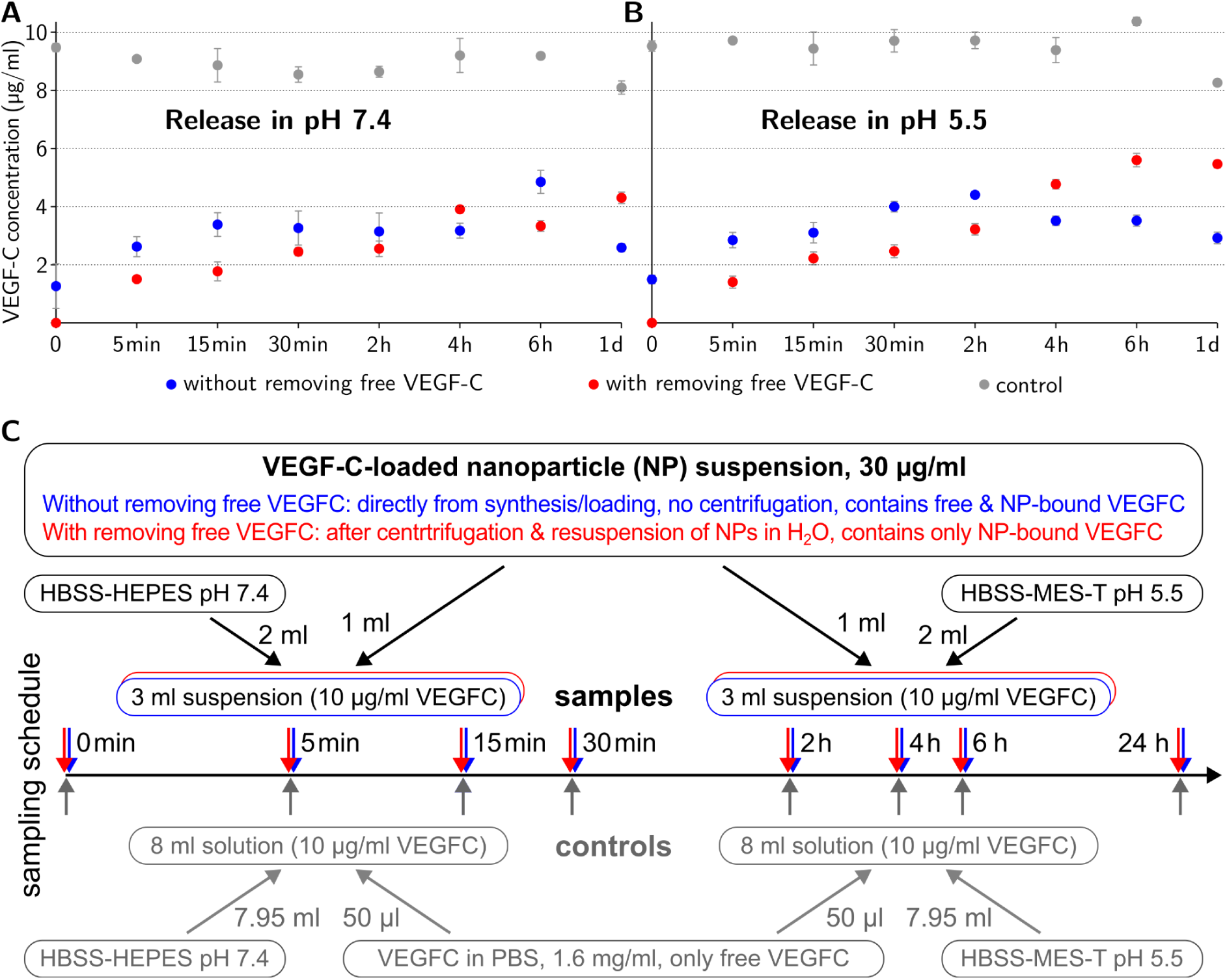
VEGFC release assays. Release assays were performed in (A) pH 7.4 and (B) pH 5.5 with and without removing the free VEGFC, as shown in the schematic (C). Since the amount of free VEGFC was not significant compared to the bound VEGFC the control is relevant for both samples, independent of whether VEGFC was removed or not.

**Supplementary Figure S3.**
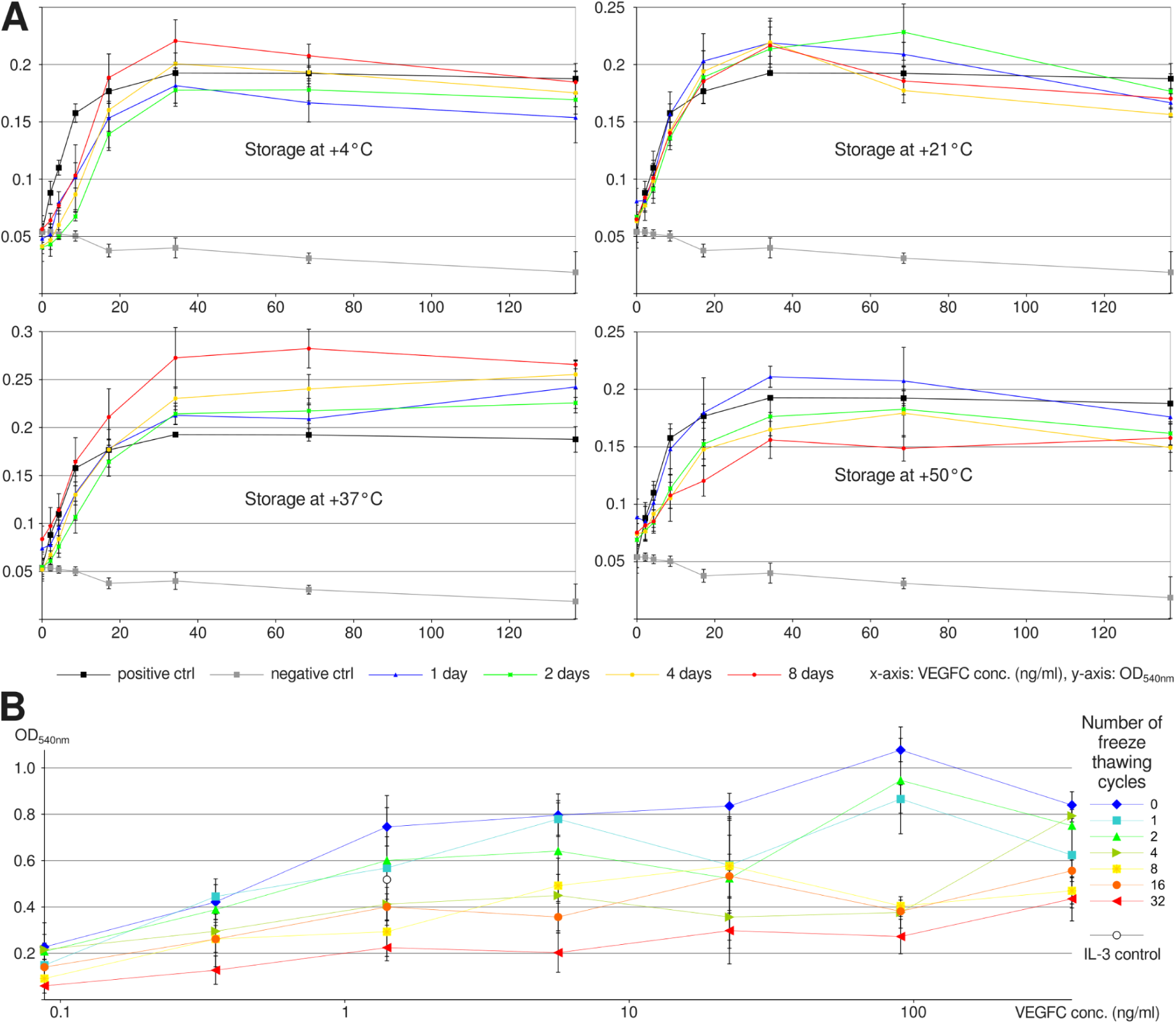
Effect of storage temperature and freeze-thaw cycles on VEGFC bioactivity. (A) Bioactivity of VEGFC as a function of storage time at different temperatures. After storing VEGFC for eight days at 37°C, we were unable to detect any loss of biological activity, and it maintains about ⅔ of it when stored for the same time at 50°C. (B) VEGFC tolerates only a limited amount of freezing-thawing events.

**Supplementary Figure S4.**
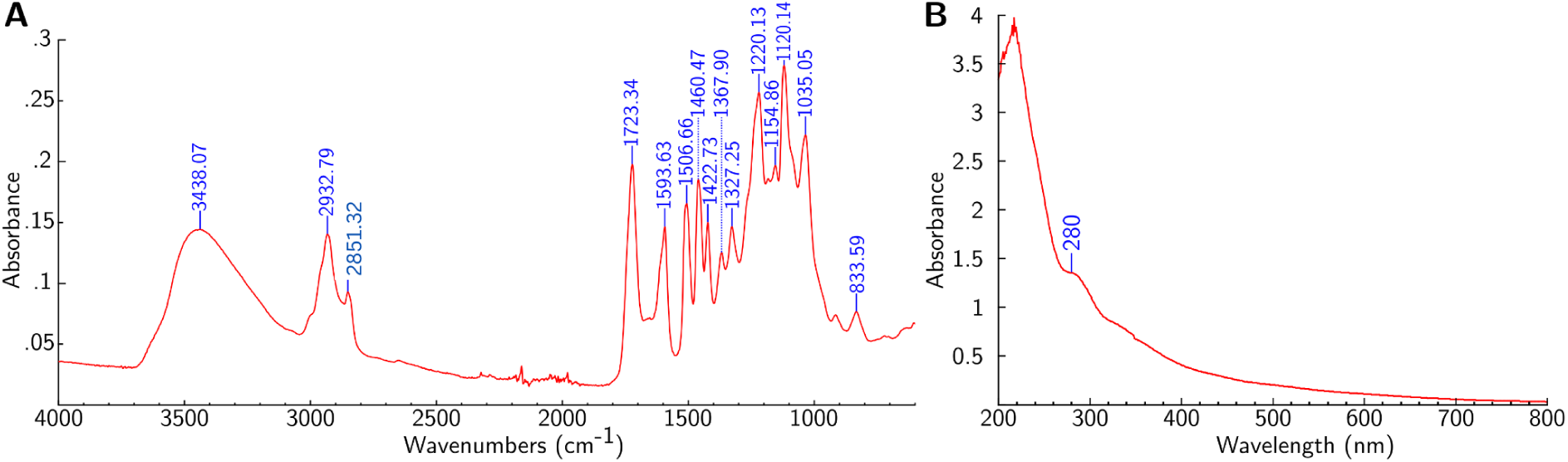
Spectroscopy characterization. (A) FTIR Spectrum of the extracted lignin. The broad absorption band at 3438.07 cm⁻¹ corresponds to the O-H stretching vibrations, typical of hydroxyl groups in lignin, indicating the extensive hydrogen bond network. The peaks at 2932.79 cm⁻¹ and 2851.32 cm⁻¹ are attributed to C-H stretching vibrations and represent stretching of C-H bond in -CH, -CH2, CH3, and -OCH3 groups, present in both aromatic and aliphatic structures.^77,78^ These peaks are common in lignin and highlight the presence of aliphatic hydrocarbons. Methoxy groups, in particular, are often abundant in lignin structures. In the fingerprint region, sharp peaks at ∼1594 cm-1 and at 1507 cm-1 are assignable to aromatic ring vibration, which are highly abundant in lignin. These peaks represent the most important and discriminating features of the lignin spectrum.^79,80^ (B) UV-Vis absorption spectrum of the extracted lignin. The 280 nm peak is attributed to a benzene ring substituted with hydroxyl or methoxy groups, indicating π-π* transitions within the aromatic structures of the lignin molecule.^44^

**Supplementary Figure S5.**
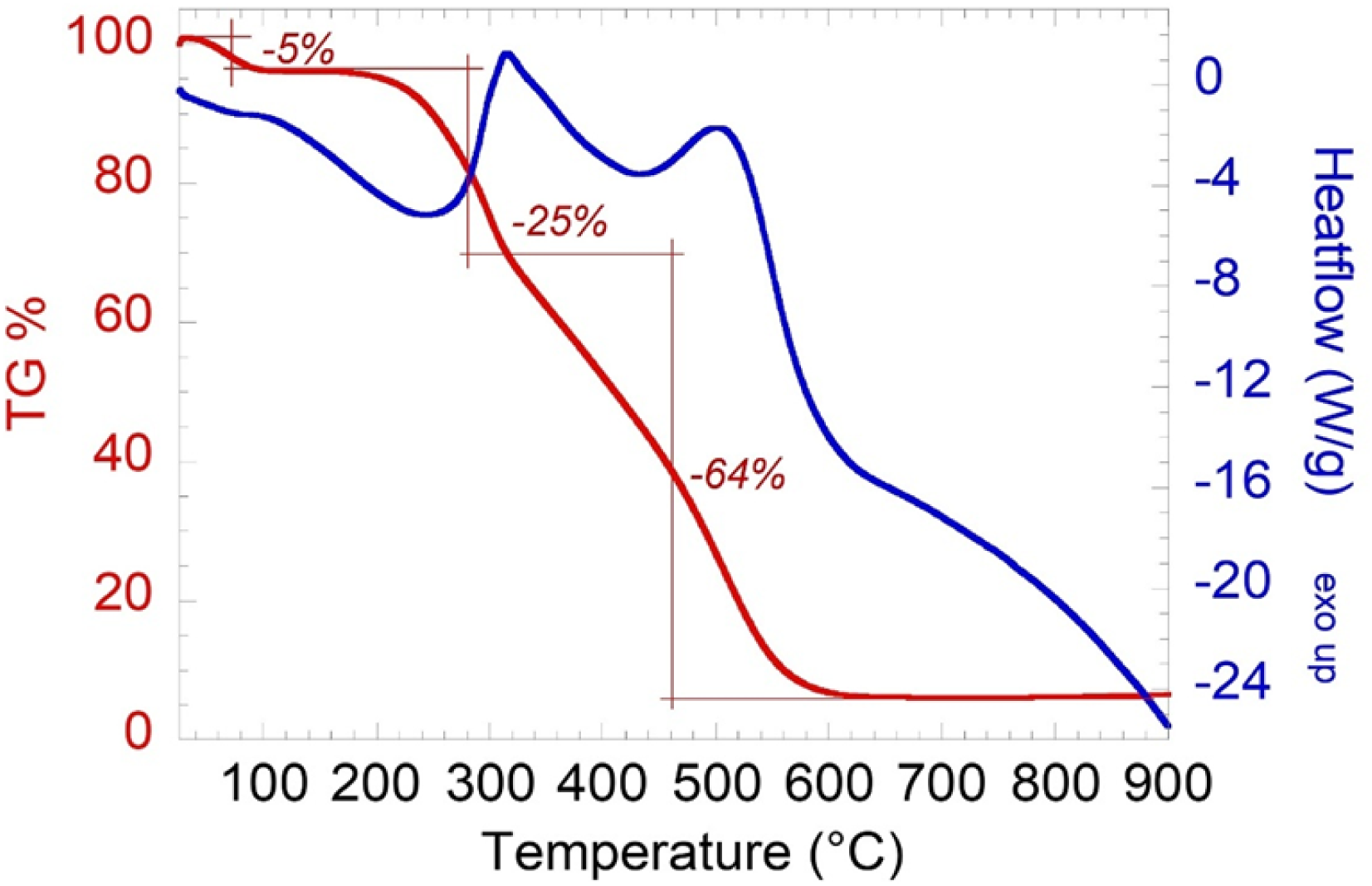
Thermal analysis of extracted lignin. The blue line shows % weight loss; the red line shows mass losses (TG); the blue graph is the DSC curve. A multi-step decomposition process is indicated by three main mass losses: a first loss of 5% weight (50-150°C), associated with moisture release due to water evaporation; a second mass loss of 25% (150-300°C), due to the thermal decomposition of hemicellulose residues linked to lignin; further decomposition, with an additional 64% weight loss (300-600°C), is marked by an exothermic heat flow signal, indicating the breakdown of stable components (see also Yang et al., 2007).^81^

**Supplementary Figure S6.**
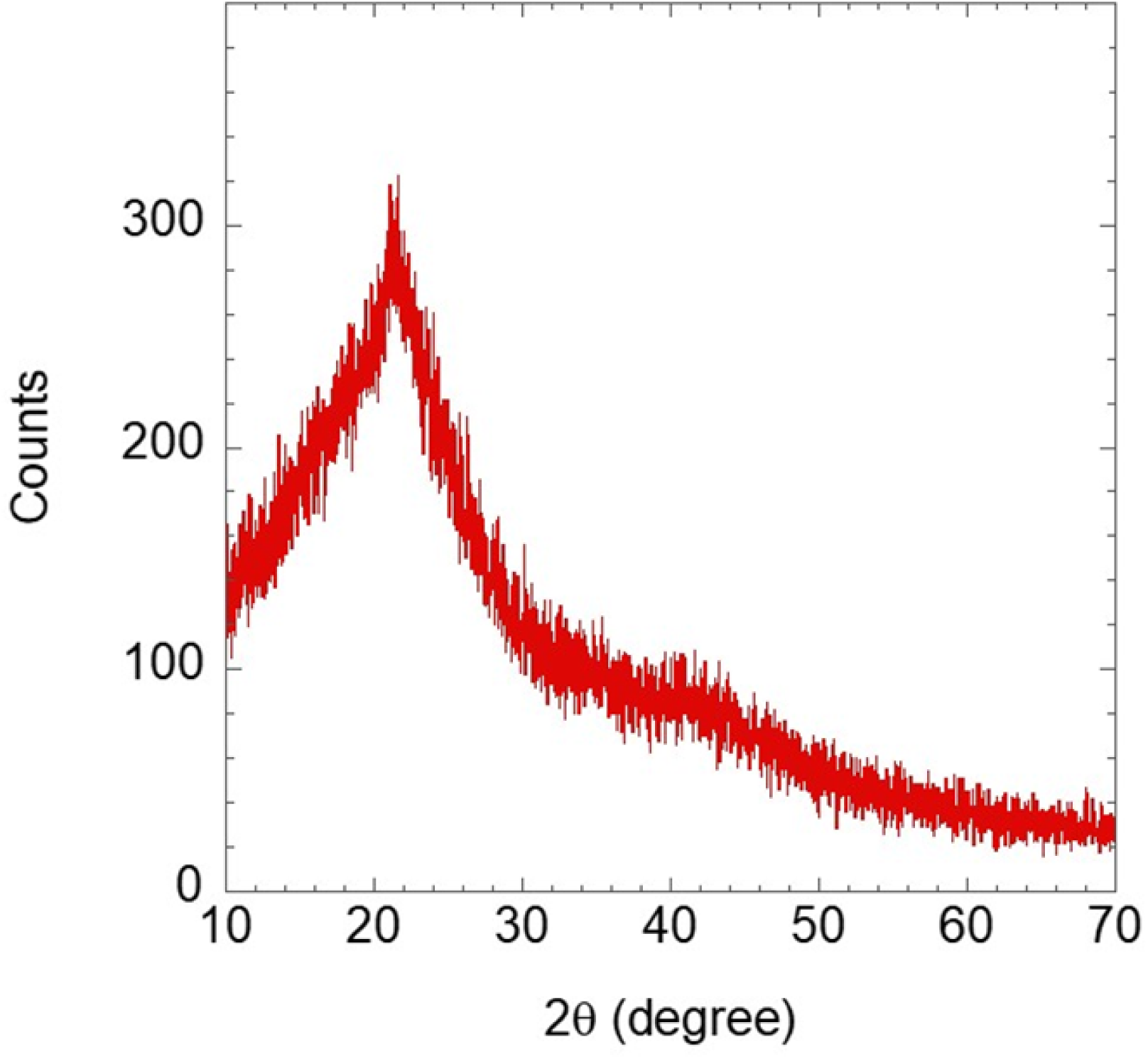
XRD graph of extracted lignin. The pattern of lignin, showing counts as a function of 2θ in the range of 10° to 70°. This pattern highlights the structural features and provides insight into the molecular organization and crystallinity of the lignin sample.

**Supplementary Figure S7.**
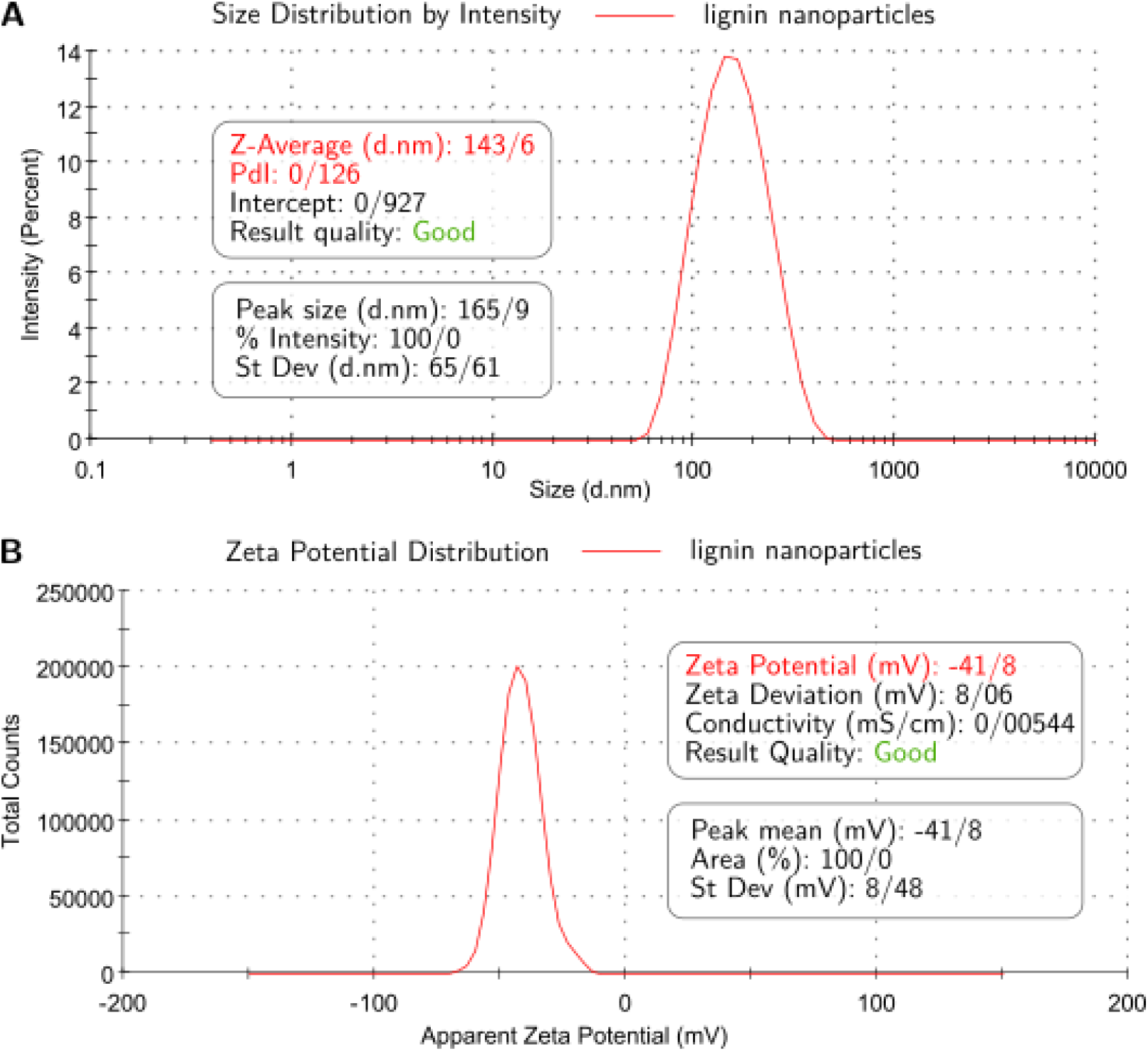
Dynamic Light Scattering (DLS) analysis of size distribution and zeta potential. (A) Particle size distribution of lignin nanoparticles, represented by intensity (%) as a function of size (nm). (B) Zeta potential distribution, showing total counts ×1000 versus apparent zeta potential (mV).

**Supplementary Figure S8.**
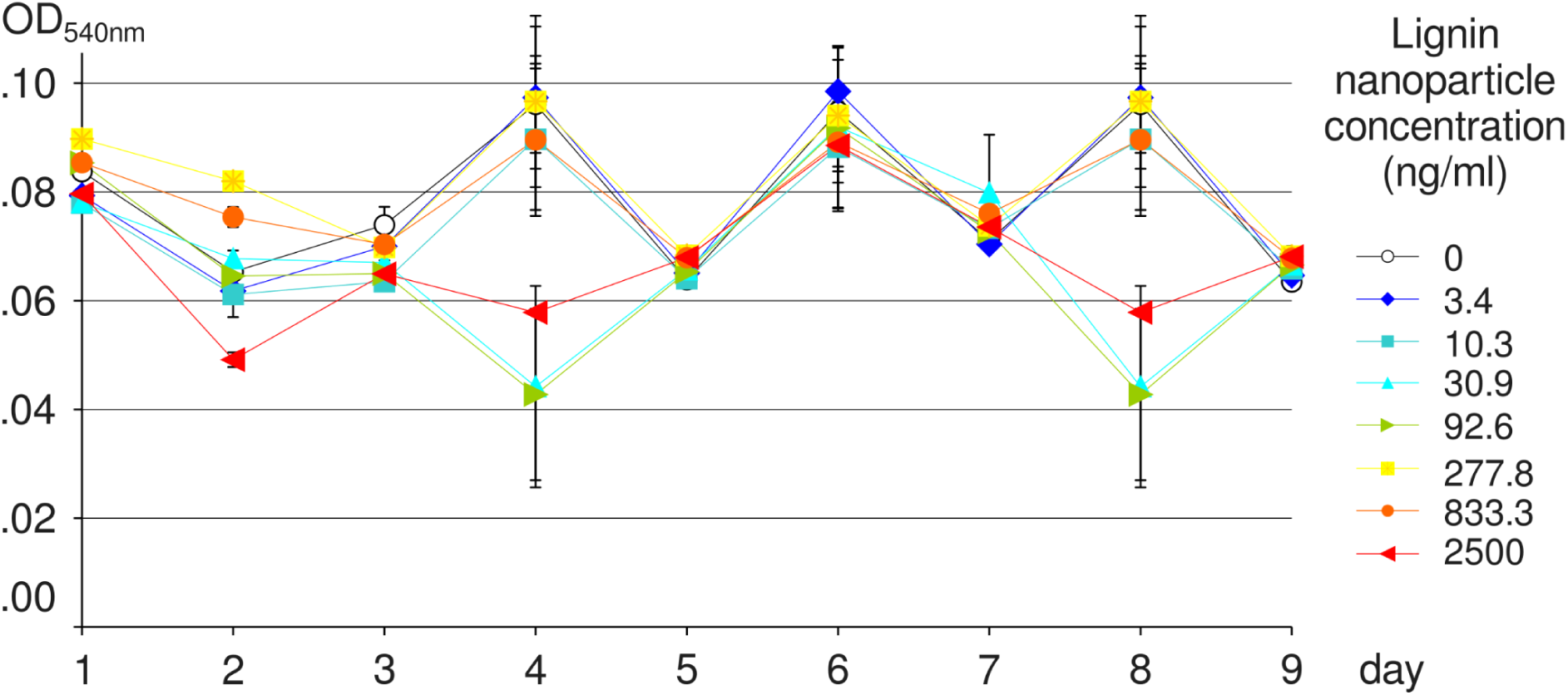
The background growth rate of Ba/F3 cells was not influenced by lignin nanoparticles, not even at high concentrations. The measurements on days 4 and 8 for the concentrations 30.9, 92.6, and 2500 ng/ml were regarded as outliers, resulting from malfunctioning of the multichannel pipette.

**Supplementary Figure S9.**
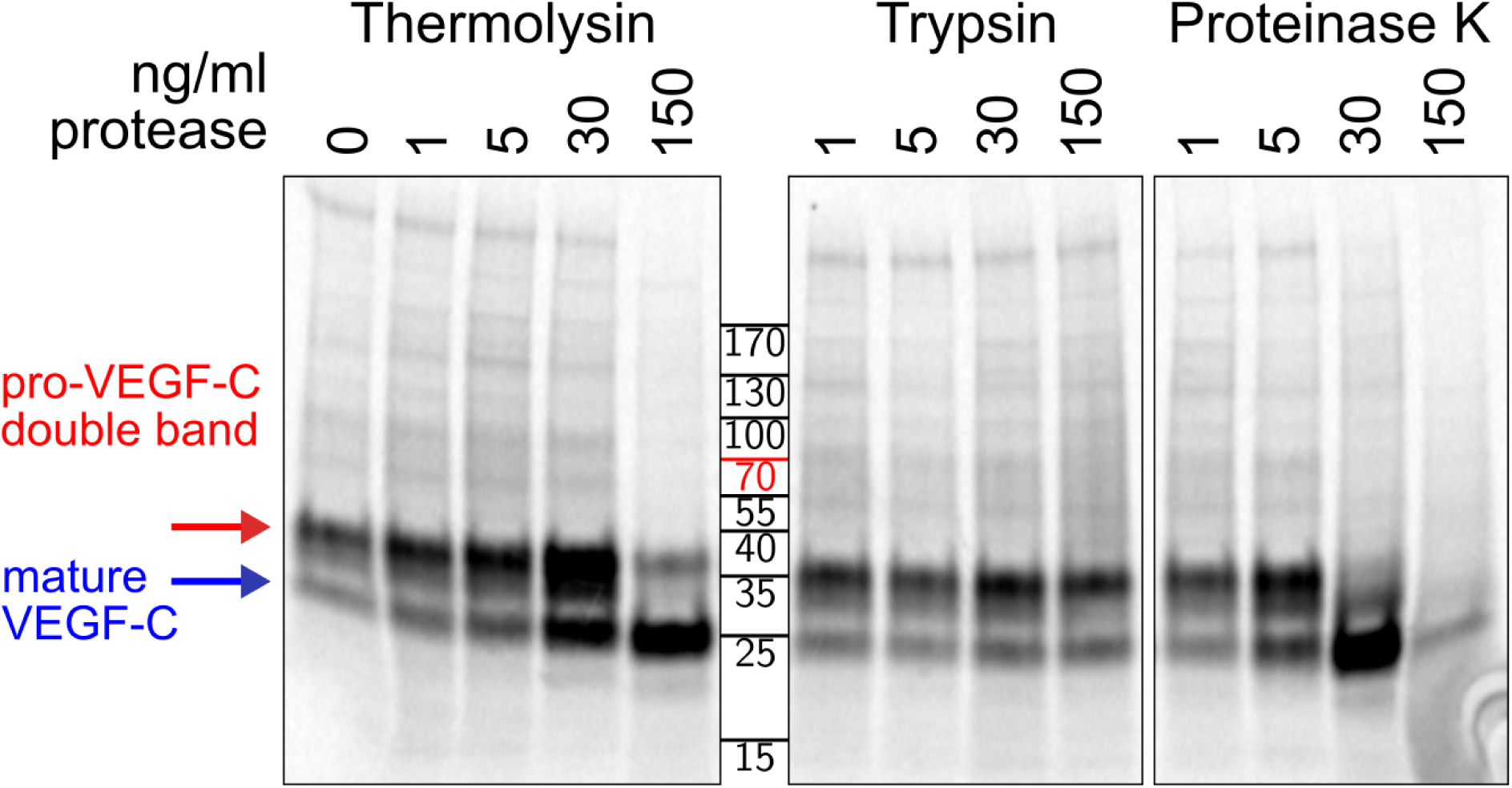
Evaluation of VEGFC degradation using gel electrophoresis. VEGFC shows relative stability against degradation by proteases in cell culture supernatant. Trypsin neither efficiently activated pro-VEGF-C nor degraded mature VEGFC, but both thermolysin and proteinase K converted pro-VEGF-C into mature VEGFC. However, the 48-hour digestion with proteinase K at 37°C degraded VEGFC nearly completely, while thermolysin only activated but didn’t degrade VEGFC. The pull-down analysis was essentially performed as previously described.^82^

**Supplementary Table T1.**
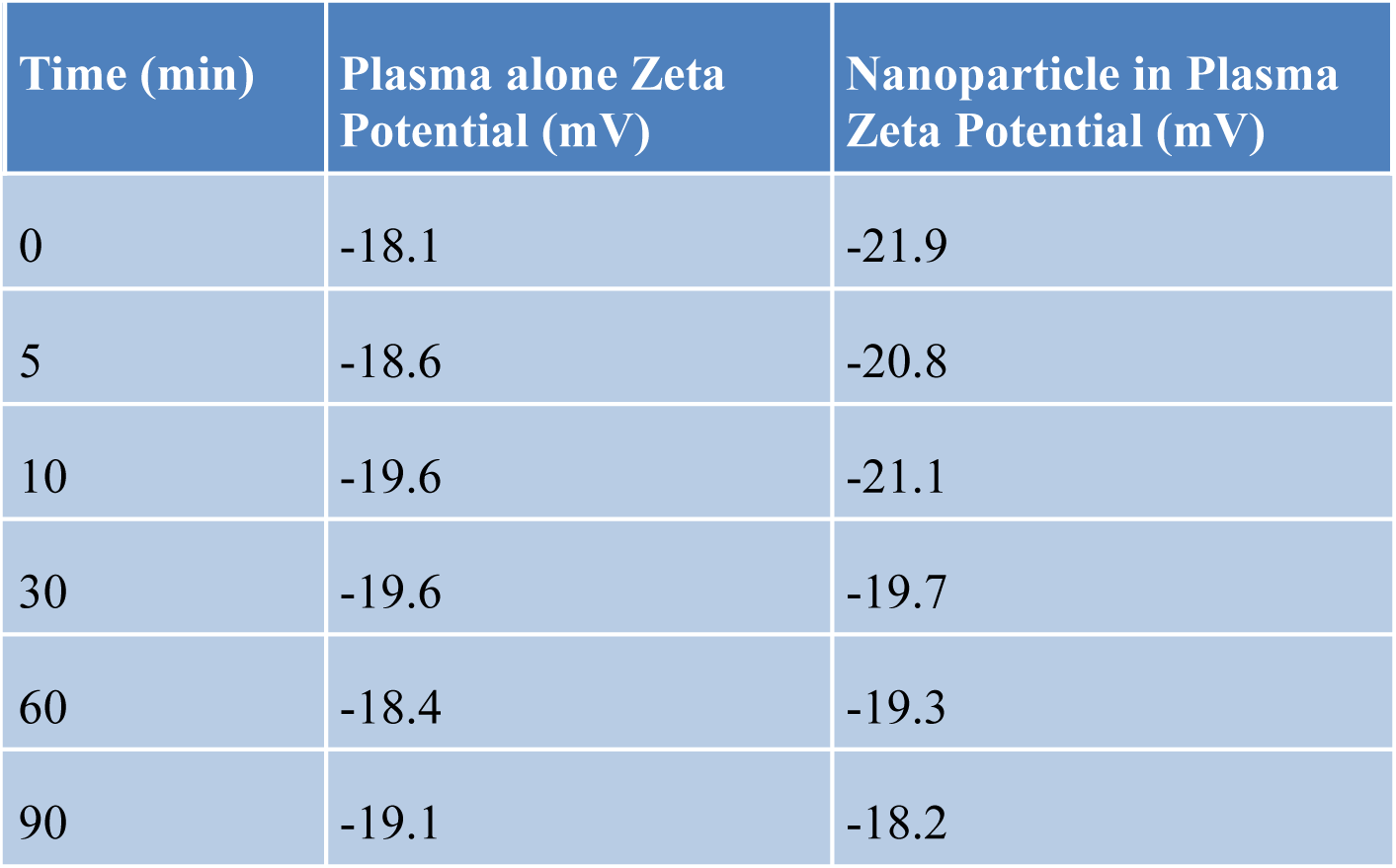
Zeta potential values (0-90 min). Zeta potential measurements over time (0–90 minutes) for LNPs in plasma and plasma alone (control) show consistent values for both conditions, indicating minimal change in surface charge throughout the observation period.

## References

(1) Wang, F.; Ouyang, D.; Zhou, Z.; Page, S. J.; Liu, D.; Zhao, X. Lignocellulosic Biomass as Sustainable Feedstock and Materials for Power Generation and Energy Storage. J. Energy Chem. 2021, 57, 247–280.

(2) Becker, J.; Wittmann, C. A Field of Dreams: Lignin Valorization into Chemicals, Materials, Fuels, and Health-Care Products. Biotechnol. Adv. 2019, 37 (6), 107360.

(3) Lizundia, E.; Sipponen, M. H.; Greca, L. G.; Balakshin, M.; Tardy, B. L.; Rojas, O. J.; Puglia, D. Multifunctional Lignin-Based Nanocomposites and Nanohybrids. Green Chem. 2021, 23 (18), 6698–6760.

(4) Ruiz-Dueñas, F. J.; Martínez, Á. T. Microbial Degradation of Lignin: How a Bulky Recalcitrant Polymer Is Efficiently Recycled in Nature and How We Can Take Advantage of This. Microb. Biotechnol. 2009, 2 (2), 164–177. 10.1111/j.1751-7915.2008.00078.x.

(5) Carmichael, E.; Ghassemieh, E.; Lyons, G. Biorefining of Lignocellulosic Feedstock and Waste Materials Using Ionic Liquid. Mater. Sci. Eng. B 2020, 262, 114741.

(6) Jiménez, L.; Pérez, A.; de la Torre, M. J.; Moral, A.; Serrano, L. Characterization of Vine Shoots, Cotton Stalks, Leucaena Leucocephala and Chamaecytisus Proliferus, and of Their Ethyleneglycol Pulps. Bioresour. Technol. 2007, 98 (18), 3487–3490.

(7) Laurichesse, S.; Avérous, L. Chemical Modification of Lignins: Towards Biobased Polymers. Prog. Polym. Sci. 2014, 39 (7), 1266–1290.

(8) Wang, H.-M.; Wang, B.; Wen, J.-L.; Yuan, T.-Q.; Sun, R.-C. Structural Characteristics of Lignin Macromolecules from Different *Eucalyptus* Species. ACS Sustain. Chem. Eng. 2017, 5 (12), 11618–11627. 10.1021/acssuschemeng.7b02970.

(9) Zhou, X.; Broadbelt, L. J.; Vinu, R. Mechanistic Understanding of Thermochemical Conversion of Polymers and Lignocellulosic Biomass. *Adv*. Chem. Eng. 2016, 49, 95–198.

(10) Yang, W.; Fortunati, E.; Gao, D.; Balestra, G. M.; Giovanale, G.; He, X.; Torre, L.; Kenny, J. M.; Puglia, D. Valorization of Acid Isolated High Yield Lignin Nanoparticles as Innovative Antioxidant/Antimicrobial Organic Materials. ACS Sustain. Chem. Eng. 2018, 6 (3), 3502–3514. 10.1021/acssuschemeng.7b03782.

(11) Wijaya, C. J.; Ismadji, S.; Gunawan, S. A Review of Lignocellulosic-Derived Nanoparticles for Drug Delivery Applications: Lignin Nanoparticles, Xylan Nanoparticles, and Cellulose Nanocrystals. Molecules 2021, 26 (3), 676.

(12) Sun, Y.; Cheng, J. Hydrolysis of Lignocellulosic Materials for Ethanol Production: A Review. Bioresour. Technol. 2002, 83 (1), 1–11.

(13) Mood, S. H.; Golfeshan, A. H.; Tabatabaei, M.; Jouzani, G. S.; Najafi, G. H.; Gholami, M.; Ardjmand, M. Lignocellulosic Biomass to Bioethanol, a Comprehensive Review with a Focus on Pretreatment. Renew. Sustain. Energy Rev. 2013, 27, 77–93.

(14) Brodeur, G.; Yau, E.; Badal, K.; Collier, J.; Ramachandran, K. B.; Ramakrishnan, S. Chemical and Physicochemical Pretreatment of Lignocellulosic Biomass: A Review. Enzyme Res. 2011, 2011, 1–17. 10.4061/2011/787532.

(15) Zhang, Z.; Harrison, M. D.; Rackemann, D. W.; Doherty, W. O.; O’Hara, I. M. Organosolv Pretreatment of Plant Biomass for Enhanced Enzymatic Saccharification. Green Chem. 2016, 18 (2), 360–381.

(16) Dai, L.; Liu, R.; Hu, L.-Q.; Zou, Z.-F.; Si, C.-L. Lignin Nanoparticle as a Novel Green Carrier for the Efficient Delivery of Resveratrol. ACS Sustain. Chem. Eng. 2017, 5 (9), 8241–8249. 10.1021/acssuschemeng.7b01903.

(17) Figueiredo, P.; Lintinen, K.; Kiriazis, A.; Hynninen, V.; Liu, Z.; Bauleth-Ramos, T.; Rahikkala, A.; Correia, A.; Kohout, T.; Sarmento, B. In Vitro Evaluation of Biodegradable Lignin-Based Nanoparticles for Drug Delivery and Enhanced Antiproliferation Effect in Cancer Cells. Biomaterials 2017, 121, 97–108.

(18) Pourmoazzen, Z.; Sadeghifar, H.; Yang, G.; Lucia, L. Cholesterol-Modified Lignin: A New Avenue for Green Nanoparticles, Meltable Materials, and Drug Delivery. Colloids Surf. B Biointerfaces 2020, 186, 110685.

(19) Zikeli, F.; Vinciguerra, V.; Sennato, S.; Scarascia Mugnozza, G.; Romagnoli, M. Preparation of Lignin Nanoparticles with Entrapped Essential Oil as a Bio-Based Biocide Delivery System. ACS Omega 2020, 5 (1), 358–368. 10.1021/acsomega.9b02793.

(20) Tortora, M.; Cavalieri, F.; Mosesso, P.; Ciaffardini, F.; Melone, F.; Crestini, C. Ultrasound Driven Assembly of Lignin into Microcapsules for Storage and Delivery of Hydrophobic Molecules. Biomacromolecules 2014, 15 (5), 1634–1643. 10.1021/bm500015j.

(21) Iravani, S.; Varma, R. S. Greener Synthesis of Lignin Nanoparticles and Their Applications. Green Chem. 2020, 22 (3), 612–636.

(22) Zhao, W.; Xiao, L.-P.; Song, G.; Sun, R.-C.; He, L.; Singh, S.; Simmons, B. A.; Cheng, G. From Lignin Subunits to Aggregates: Insights into Lignin Solubilization. Green Chem. 2017, 19 (14), 3272–3281.

(23) Lohela, M.; Bry, M.; Tammela, T.; Alitalo, K. VEGFs and Receptors Involved in Angiogenesis versus Lymphangiogenesis. Curr. Opin. Cell Biol. 2009, 21 (2), 154–165.

(24) Mandriota, S. J. Vascular Endothelial Growth Factor-C-Mediated Lymphangiogenesis Promotes Tumour Metastasis. EMBO J. 2001, 20 (4), 672–682. 10.1093/emboj/20.4.672.

(25) Karpanen, T.; Egeblad, M.; Karkkainen, M. J.; Kubo, H.; Ylä-Herttuala, S.; Jäättelä, M.; Alitalo, K. Vascular Endothelial Growth Factor C Promotes Tumor Lymphangiogenesis and Intralymphatic Tumor Growth. Cancer Res. 2001, 61 (5), 1786–1790.

(26) Sipos, B.; Klapper, W.; Kruse, M.-L.; Kalthoff, H.; Kerjaschki, D.; Klöppel, G. Expression of Lymphangiogenic Factors and Evidence of Intratumoral Lymphangiogenesis in Pancreatic Endocrine Tumors. Am. J. Pathol. 2004, 165 (4), 1187–1197.

(27) Hansel, D. E.; Rahman, A.; Hermans, J.; De Krijger, R. R.; Ashfaq, R.; Yeo, C. J.; Cameron, J. L.; Maitra, A. Liver Metastases Arising from Well-Differentiated Pancreatic Endocrine Neoplasms Demonstrate Increased VEGF-C Expression. Mod. Pathol. 2003, 16 (7), 652–659.

(28) Jha, S. K.; Rauniyar, K.; Chronowska, E.; Mattonet, K.; Maina, E. W.; Koistinen, H.; Stenman, U.-H.; Alitalo, K.; Jeltsch, M. KLK3/PSA and Cathepsin D Activate VEGF-C and VEGF-D. eLife 2019, 8, e44478. 10.7554/eLife.44478.

(29) Calvo, C.-F.; Fontaine, R. H.; Soueid, J.; Tammela, T.; Makinen, T.; Alfaro-Cervello, C.; Bonnaud, F.; Miguez, A.; Benhaim, L.; Xu, Y. Vascular Endothelial Growth Factor Receptor 3 Directly Regulates Murine Neurogenesis. Genes Dev. 2011, 25 (8), 831–844.

(30) Chen, H. I.; Poduri, A.; Numi, H.; Kivela, R.; Saharinen, P.; McKay, A. S.; Raftrey, B.; Churko, J.; Tian, X.; Zhou, B. VEGF-C and Aortic Cardiomyocytes Guide Coronary Artery Stem Development. J. Clin. Invest. 2014, 124 (11), 4899–4914.

(31) Klotz, L.; Norman, S.; Vieira, J. M.; Masters, M.; Rohling, M.; Dubé, K. N.; Bollini, S.; Matsuzaki, F.; Carr, C. A.; Riley, P. R. Cardiac Lymphatics Are Heterogeneous in Origin and Respond to Injury. Nature 2015, 522 (7554), 62–67.

(32) Ly, P.-D.; Ly, K.-N.; Phan, H.-L.; Nguyen, H. H.; Duong, V.-A.; Nguyen, H. V. Recent Advances in Surface Decoration of Nanoparticles in Drug Delivery. Front. Nanotechnol. 2024, 6, 1456939.

(33) Banerji, S.; Ni, J.; Wang, S.-X.; Clasper, S.; Su, J.; Tammi, R.; Jones, M.; Jackson, D. G. LYVE-1, a New Homologue of the CD44 Glycoprotein, Is a Lymph-Specific Receptor for Hyaluronan. J. Cell Biol. 1999, 144 (4), 789–801.

(34) Leppäpuska, I.-M.; Hartiala, P.; Suominen, S.; Suominen, E.; Kaartinen, I.; Mäki, M.; Seppänen, M.; Kiiski, J.; Viitanen, T.; Lahdenperä, O. Phase 1 Lymfactin® Study: 24-Month Efficacy and Safety Results of Combined Adenoviral VEGF-C and Lymph Node Transfer Treatment for Upper Extremity Lymphedema. J. Plast. Reconstr. Aesthet. Surg. 2022, 75 (11), 3938–3945.

(35) Zhang, L.; Gu, F.; Chan, J.; Wang, A.; Langer, R.; Farokhzad, O. Nanoparticles in Medicine: Therapeutic Applications and Developments. Clin. Pharmacol. Ther. 2008, 83 (5), 761–769. 10.1038/sj.clpt.6100400.

(36) Manara, P.; Zabaniotou, A.; Vanderghem, C.; Richel, A. Lignin Extraction from Mediterranean Agro-Wastes: Impact of Pretreatment Conditions on Lignin Chemical Structure and Thermal Degradation Behavior. Catal. Today 2014, 223, 25–34.

(37) Oh, S.-J.; Jeltsch, M. M.; Birkenhäger, R.; McCarthy, J. E. G.; Weich, H. A.; Christ, B.; Alitalo, K.; Wilting, J. VEGF and VEGF-C: Specific Induction of Angiogenesis and Lymphangiogenesis in the Differentiated Avian Chorioallantoic Membrane. Dev. Biol. 1997, 188 (1), 96–109. 10.1006/dbio.1997.8639.

(38) Karpanen, T.; Heckman, C. A.; Keskitalo, S.; Jeltsch, M.; Ollila, H.; Neufeld, G.; Tamagnone, L.; Alitalo, K. Functional Interaction of VEGF-C and VEGF-D with Neuropilin Receptors. FASEB J. 2006, 20 (9), 1462–1472. 10.1096/fj.05-5646com.

(39) Richter, A. P.; Bharti, B.; Armstrong, H. B.; Brown, J. S.; Plemmons, D.; Paunov, V. N.; Stoyanov, S. D.; Velev, O. D. Synthesis and Characterization of Biodegradable Lignin Nanoparticles with Tunable Surface Properties. Langmuir 2016, 32 (25), 6468–6477. 10.1021/acs.langmuir.6b01088.

(40) Soppimath, K. S.; Aminabhavi, T. M.; Kulkarni, A. R.; Rudzinski, W. E. Biodegradable Polymeric Nanoparticles as Drug Delivery Devices. J. Controlled Release 2001, 70 (1–2), 1–20.

(41) Zhou, Y.; Han, Y.; Li, G.; Chu, F. Effects of Lignin-Based Hollow Nanoparticle Structure on the Loading and Release Behavior of Doxorubicin. Materials 2019, 12 (10), 1694.

(42) Leppänen, V.-M.; Jeltsch, M.; Anisimov, A.; Tvorogov, D.; Aho, K.; Kalkkinen, N.; Toivanen, P.; Ylä-Herttuala, S.; Ballmer-Hofer, K.; Alitalo, K. Structural Determinants of Vascular Endothelial Growth Factor-D Receptor Binding and Specificity. Blood 2011, 117 (5), 1507–1515. 10.1182/blood-2010-08-301549.

(43) Jha, S. K.; Rauniyar, K.; Chronowska, E.; Mattonet, K.; Maina, E. W.; Koistinen, H.; Stenman, U.-H.; Alitalo, K.; Jeltsch, M. KLK3/PSA and Cathepsin D Activate VEGF-C and VEGF-D. eLife 2019, 8, e44478. 10.7554/eLife.44478.

(44) Sakakibara, A.; Sano, Y. Chemistry of Lignin. Wood Cellul. Chem. 2000, 2, 109–173.

(45) Skulcova, A.; Majova, V.; Kohutova, M.; Grosik, M.; Sima, J.; Jablonsky, M. UV/Vis Spectrometry as a Quantification Tool for Lignin Solubilized in Deep Eutectic Solvents. BioResources 2017, 12 (3), 6713–6722.

(46) Shrimal, P.; Jadeja, G.; Patel, S. A Review on Novel Methodologies for Drug Nanoparticle Preparation: Microfluidic Approach. Chem. Eng. Res. Des. 2020, 153, 728–756.

(47) Baker, A.; Kim, H.; Semple, J. L.; Dumont, D.; Shoichet, M.; Tobbia, D.; Johnston, M. Experimental Assessment of Pro-Lymphangiogenic Growth Factors in the Treatment of Post-Surgical Lymphedema Following Lymphadenectomy. Breast Cancer Res. 2010, 12 (5), R70. 10.1186/bcr2638.

(48) Stewart, D. J.; Kutryk, M. J.; Fitchett, D.; Freeman, M.; Camack, N.; Su, Y.; Della Siega, A.; Bilodeau, L.; Burton, J. R.; Proulx, G. VEGF Gene Therapy Fails to Improve Perfusion of Ischemic Myocardium in Patients with Advanced Coronary Disease: Results of the NORTHERN Trial. Mol. Ther. 2009, 17 (6), 1109–1115.

(49) Ylä-Herttuala, S.; Bridges, C.; Katz, M. G.; Korpisalo, P. Angiogenic Gene Therapy in Cardiovascular Diseases: Dream or Vision? Eur. Heart J. 2017, 38 (18), 1365–1371. 10.1093/eurheartj/ehw547.

(50) Klotz, L.; Norman, S.; Vieira, J. M.; Masters, M.; Rohling, M.; Dubé, K. N.; Bollini, S.; Matsuzaki, F.; Carr, C. A.; Riley, P. R. Cardiac Lymphatics Are Heterogeneous in Origin and Respond to Injury. Nature 2015, 522 (7554), 62–67. 10.1038/nature14483.

(51) Viitanen, T. P.; Visuri, M. T.; Hartiala, P.; Mäki, M. T.; Seppänen, M. P.; Suominen, E. A.; Saaristo, A. M. Lymphatic Vessel Function and Lymphatic Growth Factor Secretion after Microvascular Lymph Node Transfer in Lymphedema Patients. Plast. Reconstr. Surg. – Glob. Open 2013, 1 (2), 1. 10.1097/GOX.0b013e318293a532.

(52) Tammela, T.; Saaristo, A.; Holopainen, T.; Lyytikkä, J.; Kotronen, A.; Pitkonen, M.; Abo-Ramadan, U.; Ylä-Herttuala, S.; Petrova, T. V.; Alitalo, K. Therapeutic Differentiation and Maturation of Lymphatic Vessels after Lymph Node Dissection and Transplantation. Nat. Med. 2007, 13 (12), 1458–1466. 10.1038/nm1689.

(53) Leppäpuska, I.-M.; Hartiala, P.; Suominen, S.; Suominen, E.; Kaartinen, I.; Mäki, M.; Seppänen, M.; Kiiski, J.; Viitanen, T.; Lahdenperä, O.; Vuolanto, A.; Alitalo, K.; Saarikko, A. M. Phase 1 Lymfactin® Study: 24-Month Efficacy and Safety Results of Combined Adenoviral VEGF-C and Lymph Node Transfer Treatment for Upper Extremity Lymphedema. J. Plast. Reconstr. Aesthet. Surg. 2022, 75 (11), 3938–3945. 10.1016/j.bjps.2022.08.011.

(54) Ortega-Sanhueza, I.; Girard, V.; Ziegler-Devin, I.; Chapuis, H.; Brosse, N.; Valenzuela, F.; Banerjee, A.; Fuentealba, C.; Cabrera-Barjas, G.; Torres, C. Preparation and Characterization of Lignin Nanoparticles from Different Plant Sources. Polymers 2024, 16 (11), 1610.

(55) Pang, T.; Wang, G.; Sun, H.; Wang, L.; Liu, Q.; Sui, W.; Parvez, A. M.; Si, C. Lignin Fractionation for Reduced Heterogeneity in Self-Assembly Nanosizing: Toward Targeted Preparation of Uniform Lignin Nanoparticles with Small Size. ACS Sustain. Chem. Eng. 2020, 8 (24), 9174–9183. 10.1021/acssuschemeng.0c02967.

(56) Liu, Y.; Yang, G.; Jin, S.; Xu, L.; Zhao, C. Development of High-Drug-Loading Nanoparticles. ChemPlusChem 2020, 85 (9), 2143–2157. 10.1002/cplu.202000496.

(57) Yadav, P.; Yadav, A. B. Preparation and Characterization of BSA as a Model Protein Loaded Chitosan Nanoparticles for the Development of Protein-/Peptide-Based Drug Delivery System. Future J. Pharm. Sci. 2021, 7 (1), 200. 10.1186/s43094-021-00345-w.

(58) Roces, C. B.; Christensen, D.; Perrie, Y. Translating the Fabrication of Protein-Loaded Poly(Lactic-Co-Glycolic Acid) Nanoparticles from Bench to Scale-Independent Production Using Microfluidics. Drug Deliv. Transl. Res. 2020, 10 (3), 582–593. 10.1007/s13346-019-00699-y.

(59) Rinderknecht, M.; Villa, A.; Ballmer-Hofer, K.; Neri, D.; Detmar, M. Phage-Derived Fully Human Monoclonal Antibody Fragments to Human Vascular Endothelial Growth Factor-C Block Its Interaction with VEGF Receptor-2 and 3. PLoS ONE 2010, 5 (8), e11941. 10.1371/journal.pone.0011941.

(60) Ferrara, N.; Houck, K.; Jakeman, L.; Leung, D. W. Molecular and Biological Properties of the Vascular Endothelial Growth Factor Family of Proteins. Endocr. Rev. 1992, 13 (1), 18–32. 10.1210/edrv-13-1-18.

(61) Valo, H. K.; Laaksonen, P. H.; Peltonen, L. J.; Linder, M. B.; Hirvonen, J. T.; Laaksonen, T. J. Multifunctional Hydrophobin: Toward Functional Coatings for Drug Nanoparticles. ACS Nano 2010, 4 (3), 1750–1758. 10.1021/nn9017558.

(62) Valo, H.; Arola, S.; Laaksonen, P.; Torkkeli, M.; Peltonen, L.; Linder, M. B.; Serimaa, R.; Kuga, S.; Hirvonen, J.; Laaksonen, T. Drug Release from Nanoparticles Embedded in Four Different Nanofibrillar Cellulose Aerogels. Eur. J. Pharm. Sci. 2013, 50 (1), 69–77.

(63) Kumari, A.; Yadav, S. K.; Yadav, S. C. Biodegradable Polymeric Nanoparticles Based Drug Delivery Systems. Colloids Surf. B Biointerfaces 2010, 75 (1), 1–18.

(64) Cagliani, R.; Gatto, F.; Bardi, G. Protein Adsorption: A Feasible Method for Nanoparticle Functionalization? Materials 2019, 12 (12), 1991.

(65) Hadjidemetriou, M.; Kostarelos, K. Evolution of the Nanoparticle Corona. Nat. Nanotechnol. 2017, 12 (4), 288–290.

(66) Barui, A. K.; Oh, J. Y.; Jana, B.; Kim, C.; Ryu, J. Cancer-Targeted Nanomedicine: Overcoming the Barrier of the Protein Corona. Adv. Ther. 2020, 3 (1), 1900124. 10.1002/adtp.201900124.

(67) Albanese, A.; Tang, P. S.; Chan, W. C. W. The Effect of Nanoparticle Size, Shape, and Surface Chemistry on Biological Systems. Annu. Rev. Biomed. Eng. 2012, 14 (1), 1–16. 10.1146/annurev-bioeng-071811-150124.

(68) Pacifici, R. E.; Thomason, A. R. Hybrid Tyrosine Kinase/Cytokine Receptors Transmit Mitogenic Signals in Response to Ligand. J. Biol. Chem. 1994, 269 (3), 1571–1574.

(69) Achen, M. G.; Jeltsch, M.; Kukk, E.; Mäkinen, T.; Vitali, A.; Wilks, A. F.; Alitalo, K.; Stacker, S. A. Vascular Endothelial Growth Factor D (VEGF-D) Is a Ligand for the Tyrosine Kinases VEGF Receptor 2 (Flk1) and VEGF Receptor 3 (Flt4). Proc. Natl. Acad. Sci. 1998, 95 (2), 548–553.

(70) Joukov, V.; Sorsa, T.; Kumar, V.; Jeltsch, M.; Claesson-Welsh, L.; Cao, Y.; Saksela, O.; Kalkkinen, N.; Alitalo, K. Proteolytic Processing Regulates Receptor Specificity and Activity of VEGF-C. EMBO J. 1997, 16 (13), 3898–3911. 10.1093/emboj/16.13.3898.

(71) Makinen, T. Isolated Lymphatic Endothelial Cells Transduce Growth, Survival and Migratory Signals via the VEGF-C/D Receptor VEGFR-3. EMBO J. 2001, 20 (17), 4762–4773. 10.1093/emboj/20.17.4762.

(72) Imoukhuede, P. I.; Popel, A. S. Quantification and Cell-to-Cell Variation of Vascular Endothelial Growth Factor Receptors. Exp. Cell Res. 2011, 317 (7), 955–965.

(73) Ten, E.; Ling, C.; Wang, Y.; Srivastava, A.; Dempere, L. A.; Vermerris, W. Lignin Nanotubes As Vehicles for Gene Delivery into Human Cells. Biomacromolecules 2014, 15 (1), 327–338. 10.1021/bm401555p.

(74) O’Dwyer, J.; Murphy, R.; González-Vázquez, A.; Kovarova, L.; Pravda, M.; Velebny, V.; Heise, A.; Duffy, G. P.; Cryan, S. A. Translational Studies on the Potential of a VEGF Nanoparticle-Loaded Hyaluronic Acid Hydrogel. Pharmaceutics 2021, 13 (6), 779.

(75) Gousopoulos, E.; Proulx, S. T.; Bachmann, S. B.; Dieterich, L. C.; Scholl, J.; Karaman, S.; Bianchi, R.; Detmar, M. An Important Role of VEGF-C in Promoting Lymphedema Development. J. Invest. Dermatol. 2017, 137 (9), 1995–2004.

(76) Jha, S. K.; Rauniyar, K.; Karpanen, T.; Leppänen, V.-M.; Brouillard, P.; Vikkula, M.; Alitalo, K.; Jeltsch, M. Efficient Activation of the Lymphangiogenic Growth Factor VEGF-C Requires the C-Terminal Domain of VEGF-C and the N-Terminal Domain of CCBE1. Sci. Rep. 2017, 7 (1), 4916. 10.1038/s41598-017-04982-1.

(77) Boeriu, C. G.; Bravo, D.; Gosselink, R. J.; van Dam, J. E. Characterisation of Structure-Dependent Functional Properties of Lignin with Infrared Spectroscopy. Ind. Crops Prod. 2004, 20 (2), 205–218.

(78) Md Salim, R.; Asik, J.; Sarjadi, M. S. Chemical Functional Groups of Extractives, Cellulose and Lignin Extracted from Native Leucaena Leucocephala Bark. Wood Sci. Technol. 2021, 55 (2), 295–313. 10.1007/s00226-020-01258-2.

(79) Fodil Cherif, M.; Trache, D.; Brosse, N.; Benaliouche, F.; Tarchoun, A. F. Comparison of the Physicochemical Properties and Thermal Stability of Organosolv and Kraft Lignins from Hardwood and Softwood Biomass for Their Potential Valorization. Waste Biomass Valorization 2020, 11 (12), 6541–6553. 10.1007/s12649-020-00955-0.

(80) Shi, Z.; Xu, G.; Deng, J.; Dong, M.; Murugadoss, V.; Liu, C.; Shao, Q.; Wu, S.; Guo, Z. Structural Characterization of Lignin from *D*. Sinicus by FTIR and NMR Techniques. Green Chem. Lett. Rev. 2019, 12 (3), 235–243. 10.1080/17518253.2019.1627428.

(81) Yang, H.; Yan, R.; Chen, H.; Lee, D. H.; Zheng, C. Characteristics of Hemicellulose, Cellulose and Lignin Pyrolysis. Fuel 2007, 86 (12–13), 1781–1788.

(82) Jeltsch, M.; Karpanen, T.; Strandin, T.; Aho, K.; Lankinen, H.; Alitalo, K. Vascular Endothelial Growth Factor (VEGF)/VEGF-C Mosaic Molecules Reveal Specificity Determinants and Feature Novel Receptor Binding Patterns. J. Biol. Chem. 2006, 281 (17), 12187–12195.

