## Supplementary figures and images for "Biomass-derived Lignin Nanoparticles for the Sustained Delivery of Vascular Endothelial Growth Factor-C"

### Full size uncropped images

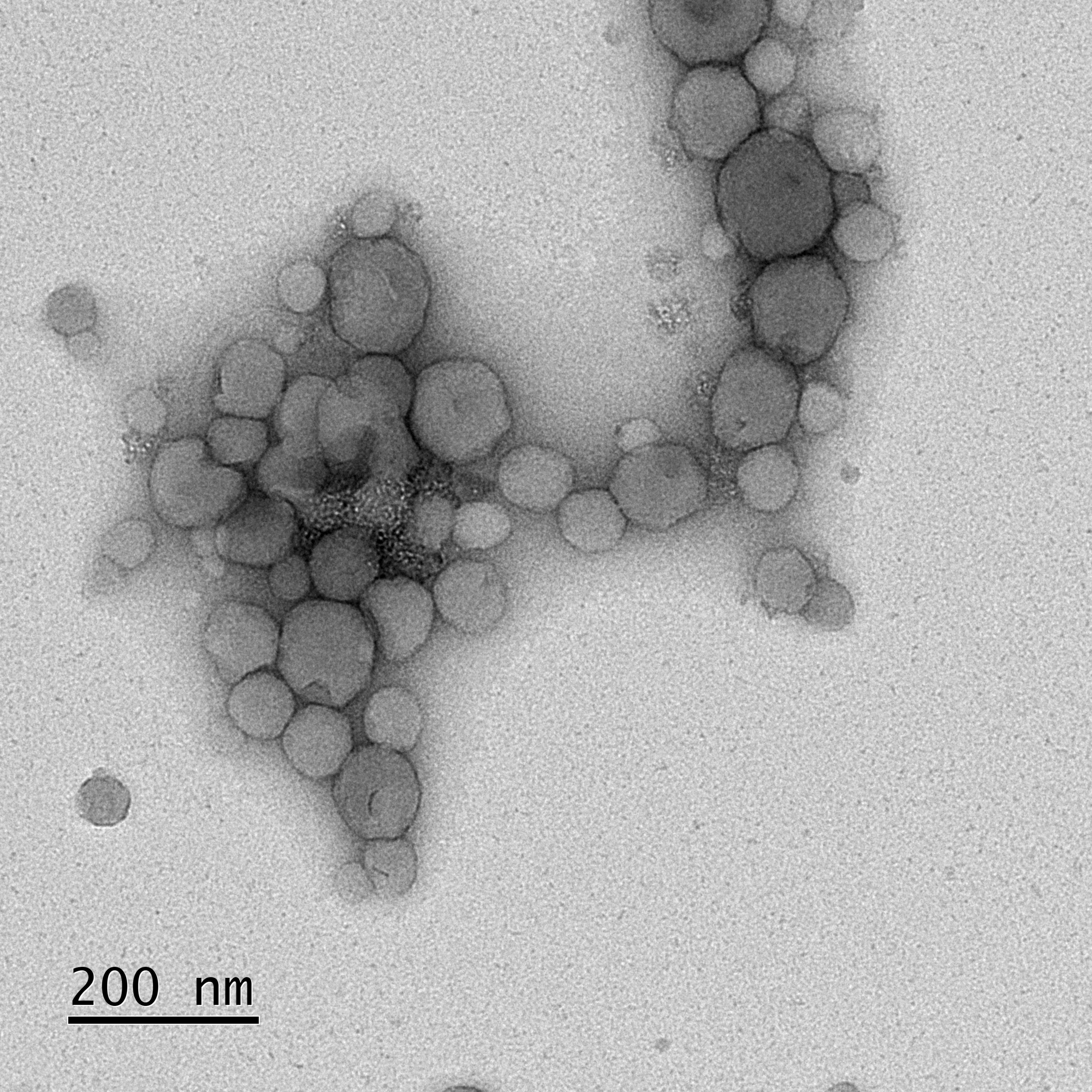

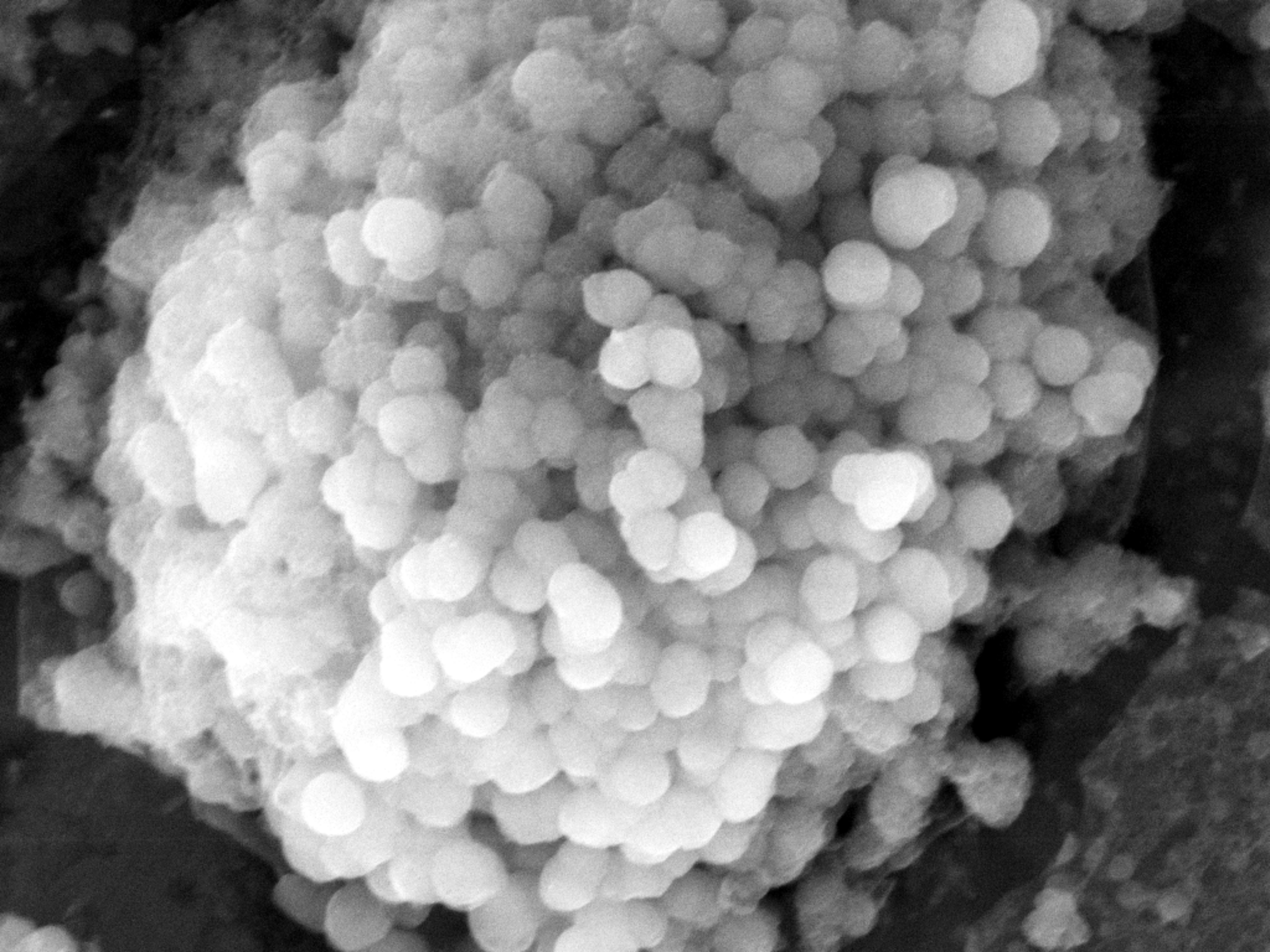

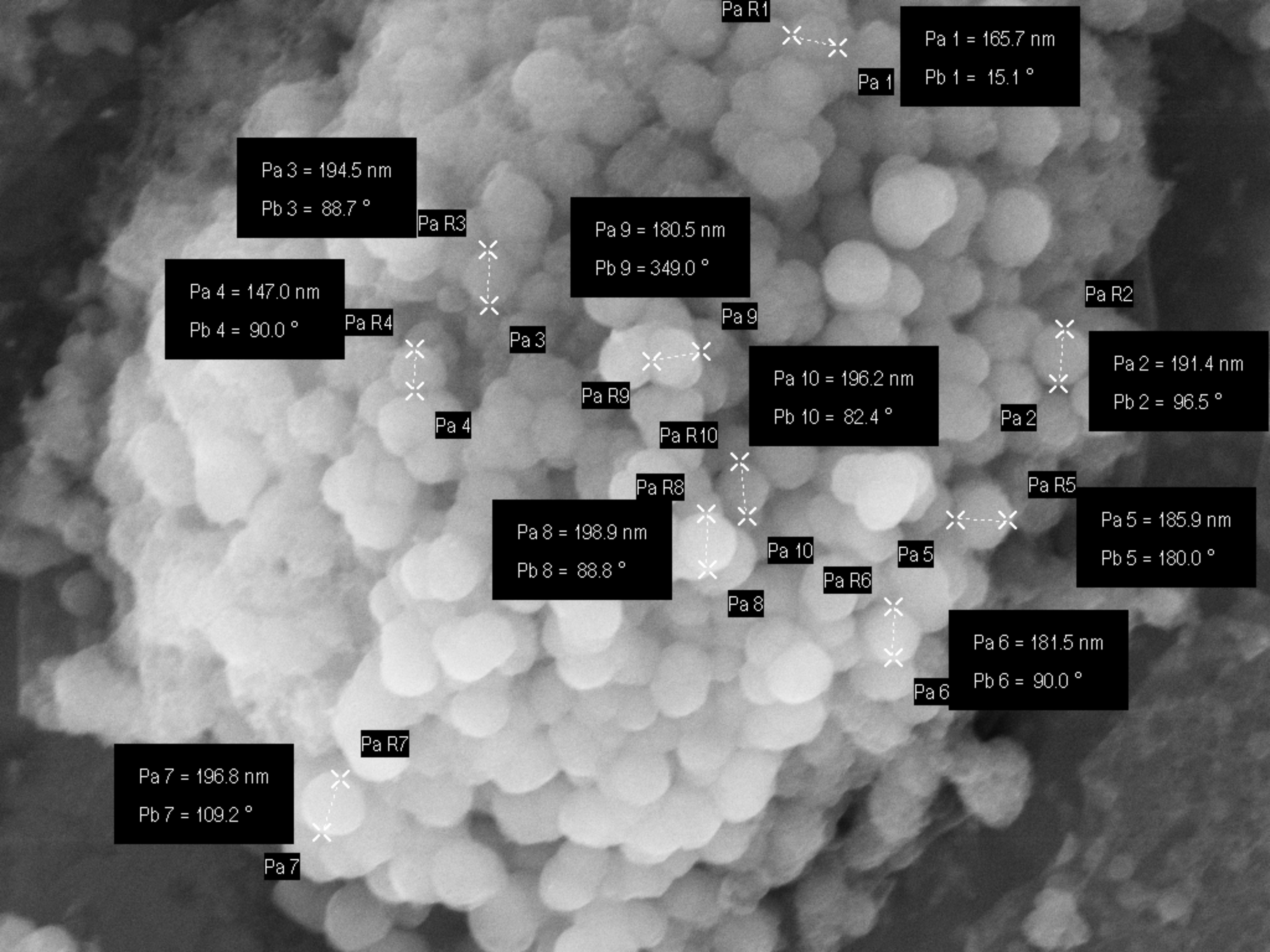

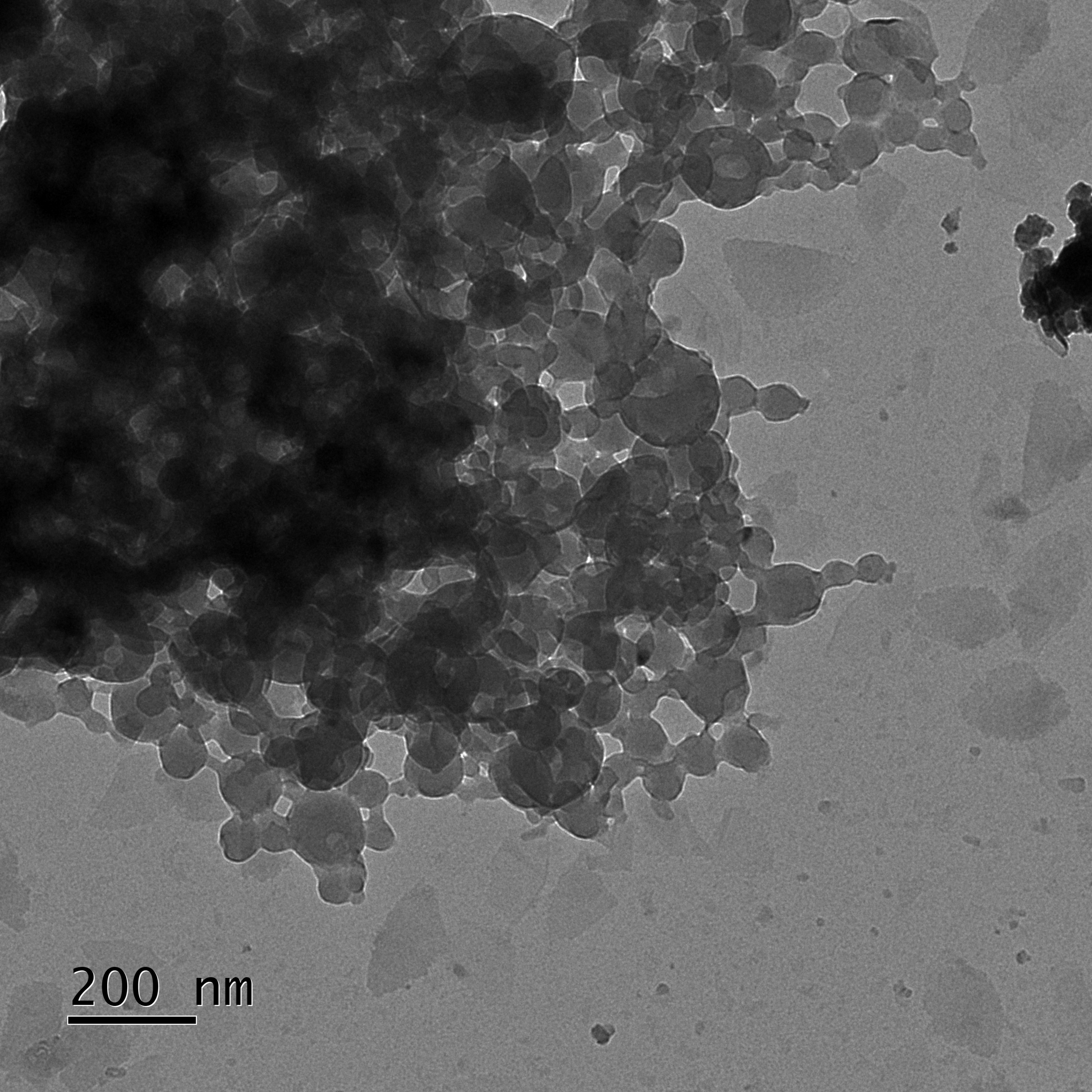

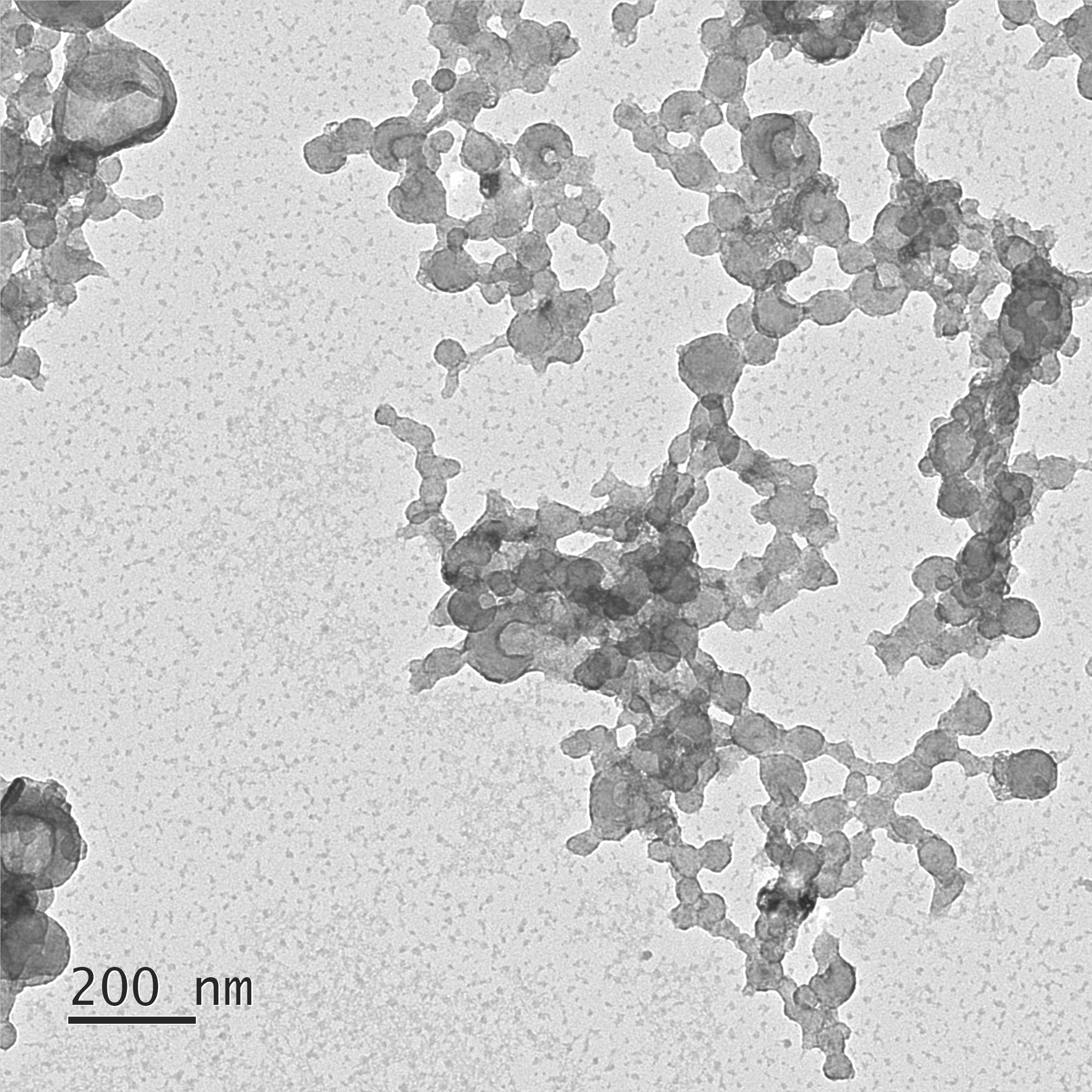

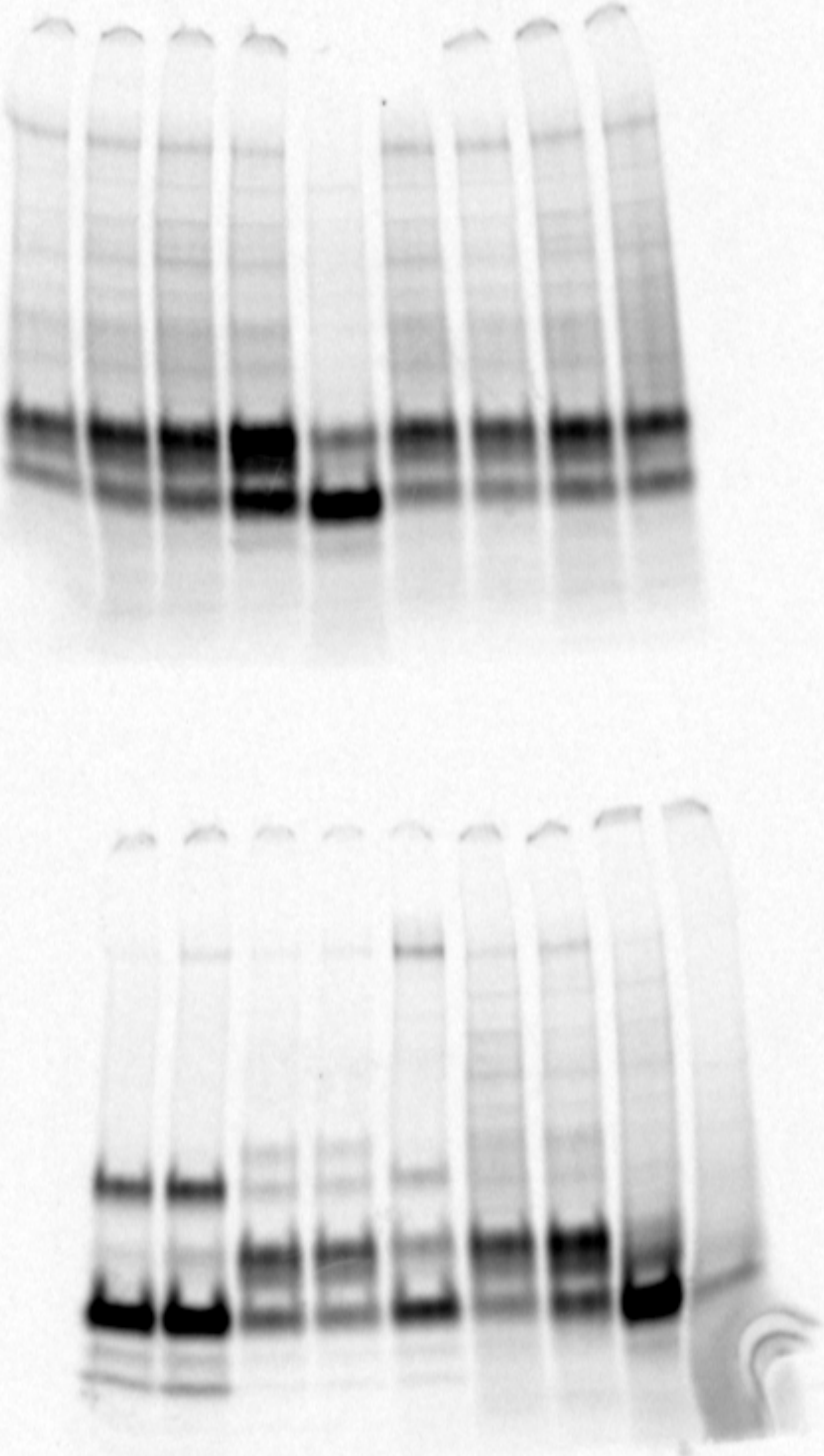

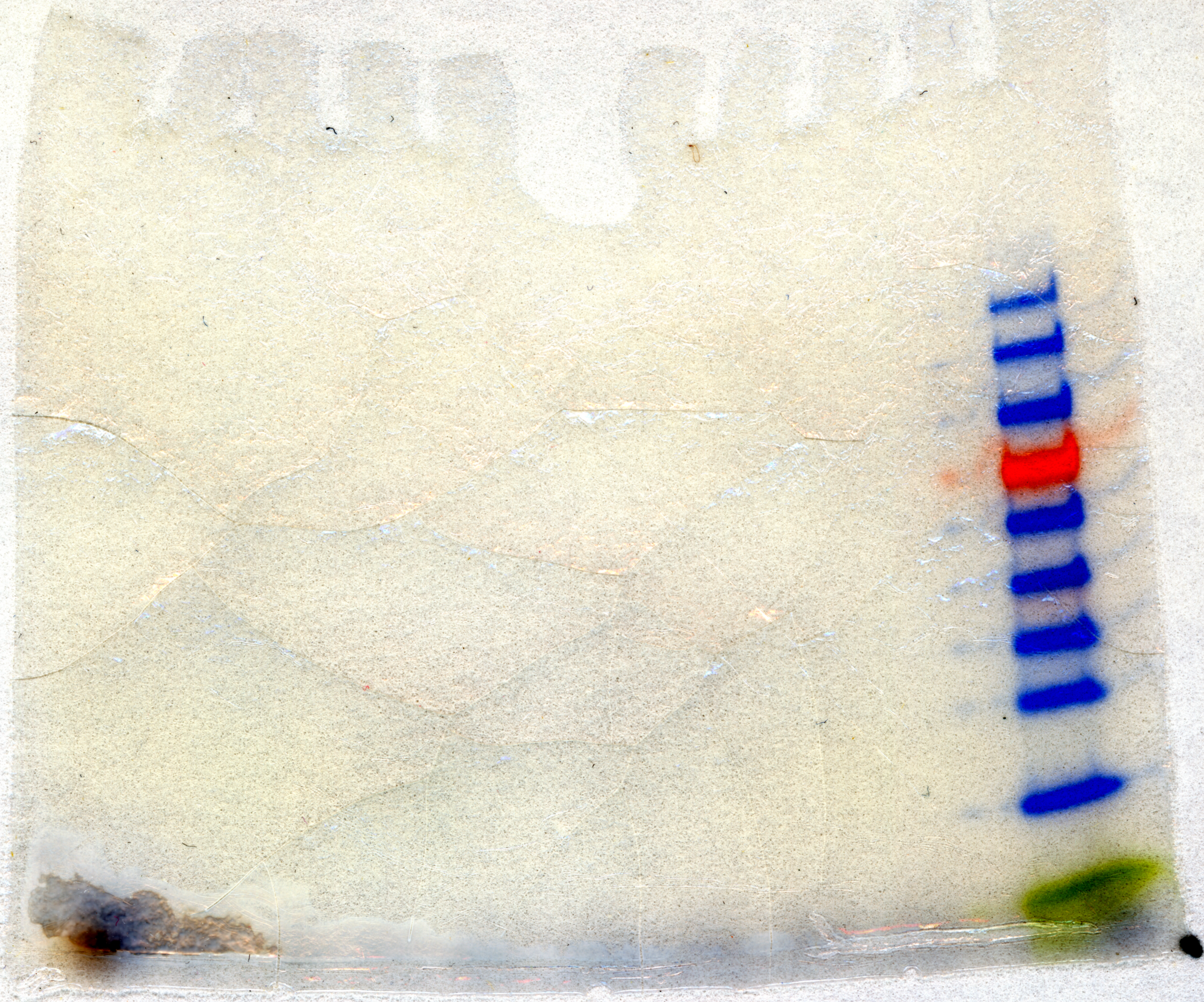

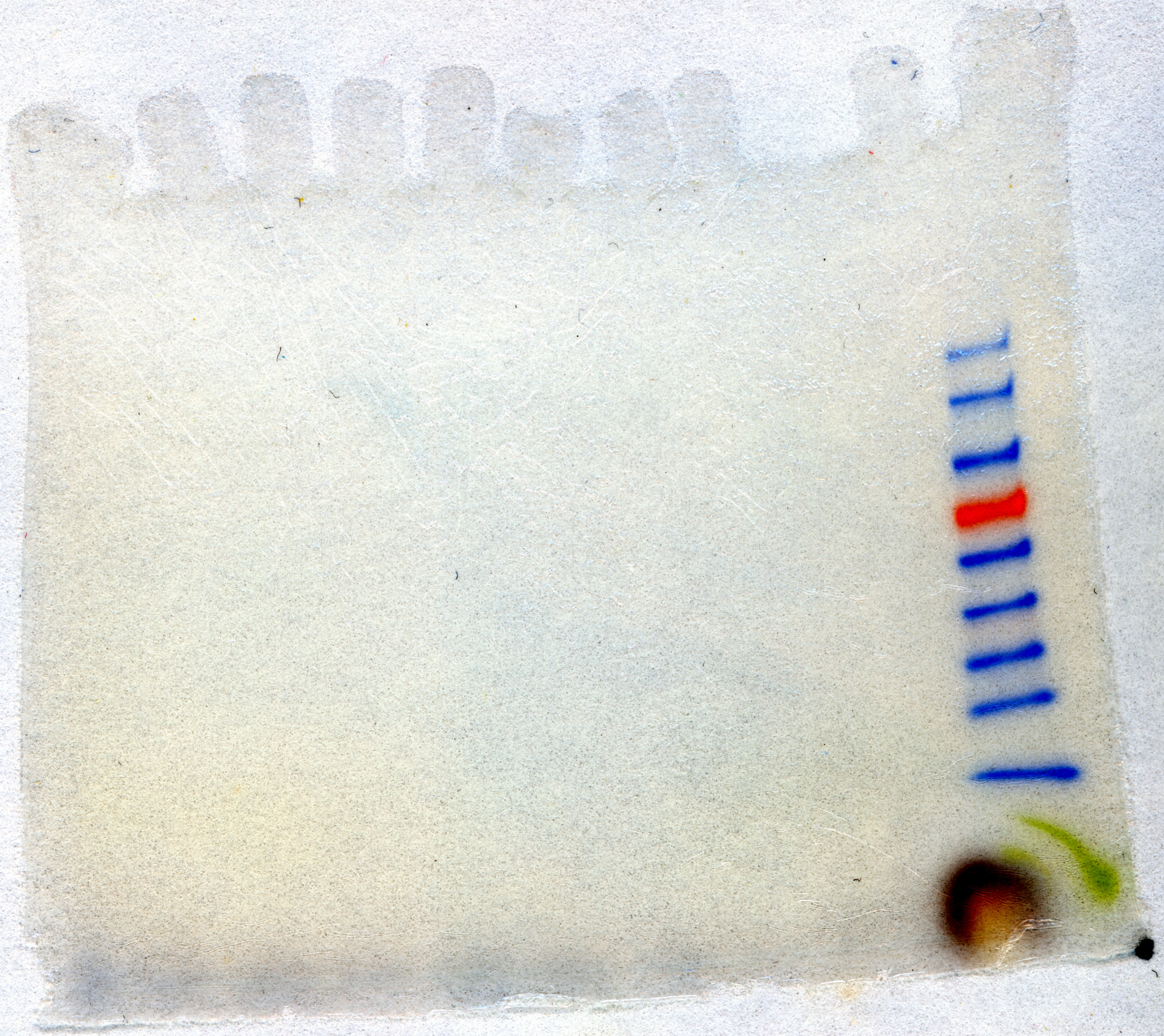
